# RNA-RNA interactions between Respiratory syncytial virus and miR-26 and miR-27 are associated with regulation of cell cycle and antiviral immunity

**DOI:** 10.1101/2023.06.05.543706

**Authors:** Sarah Ressel, Sujai Kumar, Jose Roberto Bermúdez-Barrientos, Katrina Gordon, Julia Lane, Jin Wu, Cei Abreu-Goodger, Jürgen Schwarze, Amy H. Buck

**Affiliations:** Institute of Immunology & Infection Research, School of Biological Sciences, University of Edinburgh, Edinburgh EH9 3FL, UK; Janssen Research & Development, Janssen Pharmaceutica NV, Turnhoutseweg 30, 2340, Beerse, Belgium; Institute of Ecology and Evolution, School of Biological Sciences, University of Edinburgh, Edinburgh EH9 3FL, UK; Child Life and Health, Centre for Inflammation Research, University of Edinburgh, Edinburgh EH16 4TJ, UK

**Keywords:** miRNA, respiratory syncytial virus, regulation of gene expression, pseudogenes, lncRNA, regulatory RNA, CLEAR-CLIP

## Abstract

microRNAs (miRNAs) regulate nearly all physiological processes but our understanding of exactly how they function remains incomplete, particularly in the context of viral infections. Here we adapt a biochemical method (CLEAR-CLIP) and analysis pipeline to identify targets of miRNAs in lung cells infected with Respiratory syncytial virus (RSV). We show that RSV binds directly to miR-26 and miR-27 through seed pairing and demonstrate that these miRNAs target distinct gene networks associated with cell cycle and metabolism (miR-27) and antiviral immunity (miR-26). Many of the targets are de-repressed upon infection and we show that the miR-27 targets most sensitive to miRNA inhibition are those associated with cell cycle. Finally, we demonstrate that high confidence chimeras for miR-26 and miR-27 also map to regulatory regions. We validate that a proportion of miR-27 and Argonaute 2 (AGO2) is nuclear in infected cells and identify a long non-coding RNA (lncRNA) as a miR-27 target that is linked to transcriptional regulation of nearby genes. This work expands the target networks of miR-26 and miR-27 to include direct interactions with RSV and lncRNAs and implicate these miRNAs in regulation of key genes that impact the viral life cycle associated with cell cycle, metabolism, and antiviral immunity.

## INTRODUCTION

Over the last two decades discoveries of direct RNA-RNA interactions between miRNAs and viruses have revealed novel mechanisms by which miRNAs can be regulated and by which they can exert a function. Some of the viral-miRNA interactions have been shown to increase genome stability, as first reported for the binding of miR-122 by hepatitis C virus (HCV) and more recently for miR-17 with *Pestiviruses* (1–4). Other viral-miRNA interactions inhibit the activity of the associated miRNA through sequestration from its host targets or by causing the miRNA to be degraded, as discovered in Herpesvirus saimiri (HVS) and murine cytomegalovirus (MCMV) which both use distinct viral transcripts to target miR-27 (5–8). While there is promise in blocking miRNA-virus interactions for therapeutic applications (9,10) these interactions cannot be reliably predicted.

The most reliable miRNA target prediction programs (such as TargetScan) only consider 3’ untranslated regions (3’UTRs) and coding sequences (CDS) of endogenous RNAs and are modelled based on assumptions of canonical repression and conservation (11). While these programs have enabled subsequent validation of high confidence 3’UTR targets, they are not readily applied to viral genomes. It has also been suggested that 3’UTR targeting is unlikely to explain all phenotypes associated with miRNA de-regulation (12,13), prompting the need for more systematic analyses of miRNA targets in viral infection and other diseases. In this regard, biochemical approaches have become an important complement to computational methods for miRNA target identification. Various methods have been developed to identify miRNA-mRNA interactions by UV crosslinking and immunoprecipitation (CLIP) of Argonaute (AGO) proteins followed by an intermolecular ligation step to generate chimeric sequences of miRNA and its target, including CLASH and CLEAR-CLIP (14–16). These methods have identified miRNA-target interactions in several viral families including *Flaviviridae*, however they have been refractory to the A549 cell line (4), a human lung cancer epithelial cell line that is routinely used to model gene regulation in the context of bacterial and respiratory virus infections as well as lung cancer, allergy, and asthma.

Here we first optimised the CLEAR-CLIP protocol (16) for the A549 cell line in order to build a comprehensive map of miRNA-target interactions in lung cells. We further developed a bioinformatic pipeline that maps the targets across human and viral genomes, and accounts for interactions with elements beyond CDS and 3’UTRs. We then compare targets in A549 cells that are uninfected to those infected with Respiratory syncytial virus (RSV). RSV is a 15 kb negative-sense, single-stranded RNA virus of the *Paramyxovirus* genus that has received a lot of attention recently due to surging cases after the corona virus pandemic. Despite the major clinical and economic burden in young children and the high morbidity and mortality in the elderly caused by RSV, there is still no specific therapy and only limited active immunisation available (17).

Our global CLEAR-CLIP analysis shows that miR-26 and miR-27 interact directly with RSV through seed pairing with the M gene. Analysis of host target chimeras for miR-26 and miR-27 suggest they target different sets of host genes in these cells associated with cell cycle, metabolism, and antiviral immunity. Interestingly, miR-27 host targets are statistically under-represented among the chimeric sequences upon infection, and many are sensitive to miRNA inhibition including a large network of cell cycle and metabolism genes. Through our genome analysis of miRNA-chimeric reads we further find that pseudogenes which overlap transcriptional regulatory regions are reproducible, high confidence targets. This study provides a workflow for comprehensive analysis of miRNA-target interactions in the context of respiratory virus infection, implicates miR-26 in the antiviral immune response to RSV and suggests that de-regulation of miR-27 activity during infection could impact key host genes in the viral life cycle.

## MATERIALS AND METHODS

### Cell lines and culture conditions

A549 human lung epithelial cells (CCL-185, ATCC) and human epithelial type 2 (HEp-2, CCL-23, ATCC) cells were cultured in Dulbecco’s modified eagle medium (DMEM, Sigma-Aldrich) supplemented with 10% fetal bovine serum (FBS, Gibco), 1 % L-Glutamine (L-glut, Gibco) and 1 % Penicillin/ Streptomycin (P/S, Gibco) at 37 °C and 5 % CO_2_. Cell lines were regularly checked for mycoplasma contamination by PCR with primers (forward: 5’-GGGAGCAAACAGGATTAGATACCC-3’ and reverse: 5’-TGCACCATCTGTCACTCTGTTAACCTC-3’).

### Virus and infection conditions

The human RSV, strain A2, used in this study was provided by Janssen Pharmaceuticals (Belgium). To visualise progress of viral infection a modified virus was used that had the eGFP gene introduced into the viral genome as described in Hallak et al. (18) and Guerrero-Plata et al. (19). To determine viral titre, RSV immuno-plaque assays were performed in HEp-2 cells using 2-fold serial dilutions of the virus samples in DMEM. Cells were incubated with the virus dilution for 2 h before DMEM supplemented with 10 % FBS, 1 % L-glut, 1 % P/S was added, and the cells were incubated for further 22 h. The cells were then probed with 1:200 biotinylated RSV (Biorad) antibody and stained with 1:500 ExtrAvidin Peroxidase, with visualization of plaques by the addition of 3-Amino-9-EthylCarbazole (AEC) substrate. For infection assays in A549, the cells were seeded 24 h prior to infection and infected with RSV at a multiplicity of infection (MOI) of 0.1 in serum free DMEM (+1 % L-glut) for 1 h. Subsequently the cells were washed with serum free DMEM and incubated for a total of 48 h from the start of the infection in complete media (DMEM + 10 % FBS, 1 % L-glut) unless stated otherwise.

## CLEAR-CLIP

The protocol described here was modified from the originally published CLEAR-CLIP(16) and modifications regarding buffers were guided by a variation of the protocol published by Bjerke and Yi (20). We further adapted the protocol to include a custom 3’ adapter from TriLink (developed here) and adaptation to remove need for membrane transfer (developed here). The modified protocol is described below.

### Sample preparation

A549 cells were grown in 245 mm square dishes. One day post-seeding cells were at about 80-90% confluency and either RSV-infected at an MOI of 0.1 or Mock-infected for 48 h. The plates were then washed with PBS and UV crosslinked (245 nm) on ice, covered with ice-cold PBS and UV irradiated at 200 mJ/cm^2^ (or compared to 400 mJ/cm^2^ or no crosslinking as detailed). The cells were harvested by scraping and immediately lysed in CLEAR-CLIP lysis buffer (50 mM Tris/HCl pH 7.5, 100 mM NaCl, 1 mM MgCl_2_, 0.1 mM CaCl_2_, 1 % Igepal, 0.5 % sodium deoxycholate, 0.1 % SDS, protease inhibitors (Roche)). After 10 min incubation on ice, lysates were DNase treated with 30 µL RQ1 RNase-free DNase and RNase treated with RNace-IT (Agilent) at a final concentration of 1:25,000 for 5 min at 37 °C with shaking. Lysates were cleared by centrifugation and 200 U RNasin RNase inhibitor (Promega) was added.

### Immunoprecipitation

Protein G Dynabeads (Life Technologies) were prepared using 100 µL of beads coupled to 10 µg of AGO2 antibody (clone 11A9, Merck) or IgG control antibody (Rat IgG2a kappa Isotype Control (eBR2a), Invitrogen) per sample. Beads were equilibrated in lysis buffer before the lysate was added to the beads and incubated rotating at 4 °C overnight. The beads were then washed as followed.

- 2x CLEAR-CLIP lysis buffer (1x the beads were directly resuspended in buffer, 1× 3 min rotation at 4 °C)
- 3x HS-CLEAR (50 mM Tris-HCl pH 7.4, 1 M NaCl, 1 mM EDTA, 1 % Igepal, 0.5 % sodium deoxycholate, 0.1 % SDS) for 3 min rotating at 4 °C each,
- 2x PNK+Tween (20 mM Tris-HCl pH 7.4, 10 mM MgCl_2_, 0.2 % Tween-20)

All following reactions included 80 U RNase inhibitors and incubations were carried out with intermittent shaking.

### 5’ phosphorylation, intermolecular ligation and 3’ adapter ligation

The samples were then treated with 40 U of T4 polynucleotide kinase (PNK, 3’ phosphatase minus, NEB) in a total reaction volume of 80 µL, including 10 mM ATP for 2.5 h at 20 °C to add a 5’ phosphate group for the intermolecular ligation. Samples were subsequently washed 3x in PNK+Tween buffer. After the last wash, beads were flash centrifuged to remove residual buffer. The intermolecular RNA-RNA ligation was performed in a total volume of 100 µL using 62.5 U T4 RNA ligase 1 (NEB), 100 mM ATP, 15 % PEG-8000 at 16 °C overnight. The next day more ligase (25 U) and ATP (100 mM) were added to the reaction followed by further incubation for 5 h at 16 °C. The samples were washed 2x with CLEAR lysis buffer, followed by one wash in PNK/EDTA/EGTA buffer (50 mM Tris-HCl pH 7.4, 10 mM EDTA, 10 mM EGTA, 0.5 % Igepal) and two PNK+Tween washes. The samples were then treated with 8 U TSAP for 45 min at 20 °C and washed 2x in PNK+Tween buffer. A modified 3’ sequencing adapter (3’ CleanTag-IRDye800CW, TriLink, see Table S9, 1 µM) was ligated overnight at 16 °C in the dark using 800 U T4 RNA ligase 2 (truncated K227Q, NEB), 10 % PEG-8000 in a total reaction volume of 80 µL.

### Elution from beads and SDS-PAGE purification of AGO2-RNA complexes

Beads were washed 2x with PNK+Tween and resuspended in elution buffer (15 μL 4x NuPAGE LDS sample buffer (Thermo Scientific), 45 μL PNK buffer (50 mM Tris-HCl pH 7.4, 50 mM NaCl, 10 mM MgCl2, 0.5 % TritonX-100, 5 mM β-Mercaptoethanol), 3 μL 1 M DTT). Samples were eluted from the beads at 70 °C with constant shaking. Eluate, containing AGO2-RNA complexes were run on a 4-12 % NuPAGE Bis-Tris protein gel for 150 min at 120 V in MOPS SDS running buffer (Thermo Scientific). Gels were run on ice to prevent RNA degradation and in the dark to protect the 3’ adapter fluorophore. Fluorescent RNA was visualised using the Odyssey CLx scanner (Li-cor). The regions corresponding to 80-115 kDa or 115-185 kDa (see Table S1) were excised from the gel using a scalpel. Gel pieces were flash frozen on dry ice for 5 min and fragmented by centrifugation (6 min, 6,000 rpm) through a hole in a 0.5 mL tube into a 1.5 mL tube.

### Protein digest, RNA extraction and sequencing

The shredded gel pieces were treated with 200 μg Proteinase K in 400 μL Proteinase K buffer (50 mM Tris-HCl pH 7.5, 50 mM NaCl, 0.1 % TritonX-100, 10 mM Imidazole, 1 % SDS, 5 mM EDTA) and 1 μL RNasin RNase inhibitor (Promega). The mix was incubated for 2 h at 55 °C with constant shaking. To extract the RNA an equal volume of Acid-phenol:chloroform (Ambion) was added, and the extracted RNA was precipitated over the weekend. Precipitated RNA was washed with 70 % ethanol, air dried on ice and resuspended in 2.5 μL RNase-free H_2_O and 2.5 μL buffer 1 (CleanTag Small RNA library kit). Small RNA libraries were generated using the CleanTag Small RNA library kit (TriLink) following manufacturer’s instructions with the following modifications. As the 3’ ligation was already complete, library preparation was continued at the 5’ adapter ligation step. The adapter was diluted 1:10, libraries were amplified using 20 PCR cycles and a size range of 160-238 bp was purified from the TBE gel (to avoid the miRNA only band). Samples were then sequenced by Edinburgh Genomics on Illumina NovaSeq using 100 single-end or by Genewiz (Azenta Life Sciences, Germany) using 150 nt paired-end as indicated in Table S1.

### Transfection of miRNA mimics/inhibitors, siRNAs and GapmeRs

Synthetic miRNA mimic and inhibitors as well as controls were obtained from Dharmacon, siRNAs were obtained from ThermoFisher and GapmeRs were obtained from Qiagen (see Table S9). A549 cells were reverse transfected using 5 or 10 nM of the respective mimic/inhibitor/siRNA/GapmeRs in 0.3 % Lipofectamine 2000 (Invitrogen). These were pre-incubated with Lipofectamine in Opti-MEM I (Gibco) for 20 min and subsequently added to empty wells of either a 96 well or 6 well plate. The cell mix was added to the transfection mix in antibiotic free DMEM, supplemented with 10 % FBS and 1 % L-Glut. Transfected cells were incubated for 24 h at 37 °C and 5 % CO_2_ before cell viability was measured using the CellTiter-Blue reagent (Promega) according to manufacturer’s instructions and cells were infected with RSV.

### RNA purification and sequencing

Total RNA was extracted from cells using miRNeasy mini kit (Qiagen) according to manufacturer’s instructions. If RNA was being used for total RNA-seq, the samples were on-column DNase-treated according to manufacturer’s instructions (Qiagen). RNA concentrations were determined using the Qubit RNA HS assay kit (Invitrogen). RNA quality was assessed using Bioanalyzer Pico chips (Agilent).

For total RNA-seq, libraries were prepared by the Genetics Core in Edinburgh using the NEBNext Ultra 2 Directional RNA library prep kit for Illumina and the NEBNext rRNA Depletion kit (Human/Mouse/Rat) according to the provided protocol. Sequencing was performed on the NextSeq 2000 platform (Illumina) using NextSeq 1000/2000 P3 Reagents (200 Cycles) v3. For small RNA-seq, 1 µg of RNA was treated with 5’ polyphosphatase (Cambio) according to kit instructions, followed by ethanol precipitation of the RNA. Libraries were generated from 100 ng of RNA using the CleanTag Small RNA library kit (TriLink) according to the manufacture’s instruction with the following modifications. Adapters were diluted 1:2, libraries were amplified using 15 PCR cycles and a size range of 147-238 bp was purified from the TBE gel. Libraries were sequenced on the Illumina NovaSeq (Edinburgh Genomics) using 100-base singe end.

### Reverse transcription and quantitative PCR

Extracted RNA was reverse transcribed using the QuantiTect RT kit (Qiagen, for total RNA) or miRCURY RT kit (Qiagen, for small RNAs) according to manufacturer’s instructions. The resulting cDNA was diluted as per kit instructions and quantified with specific primers (Table S9) using either QuantiNova SYBR green assays (Qiagen, for cDNA from total RNA) or miRCURY SYBR green assays (Qiagen, for cDNA from small RNAs). Small RNA miRCURY qPCR primer assays were purchased from Qiagen. Primers for the PSG lncRNA were designed to match the PSG lncRNA perfectly but have a mismatch at the 3’ end to the parental ZFAND5 gene. Mismatches located in the 3’ end were shown to impair PCR amplification (21–23) and allow for differentiation between PSG lncRNA and parental gene. ZFAND5 parental gene primers are located in exon 6 (forward primer) and exon 7 (reverse primer), which according to our data are not expressed at the PSG locus and should only amplify mature mRNA. Data were analysed using the 2^-ΔΔCt^ method (24).

### Nuclear fractionation

The nuclear fractionation protocol was adapted from Burke and Sullivan (25). Briefly, cells were harvested and washed with PBS before being lysed in CSKT buffer (10 mM PIPES (pH 6.8), 100 mM NaCl, 300 mM sucrose, 3 mM MgCl_2_, 1 mM EDTA, 1 mM DTT, 0.5 % Igepal, protease inhibitors (Roche)) on ice for 10 min. Nuclei were pelleted by centrifugation and cytoplasmic fraction (supernatant) was removed. Cytoplasmic fraction was either mixed with an equal amount of SDS lysis buffer (1 % SDS, 2 % β-Mercaptoethanol), boiled at 95°C for 10 min and used for Western blot analysis or RNA was ethanol precipitated for subsequent RNA extraction with the miRNeasy mini kit. Nuclei were washed in CSKT buffer and either lysed directly in SDS lysis buffer for Western blot analysis as described above or Qiazol for RNA extraction. To generate whole cell lysates, harvested cells were directly lysed in SDS lysis buffer/ Qiazol. Equal volumes were maintained for cytoplasmic and nuclear fractions. To test chromatin association of AGO2, the washed nuclei were lysed in CLEAR-CLIP lysis buffer for 20 min. Nuclear soluble fraction (supernatant) was separated from the insoluble fraction (pellet, containing chromatin) by centrifugation. The insoluble fraction was then sonicated 3x 10 s with 30 s on ice in between, followed by treatment with 30 µL DNase RQ1 for 5 min at 37 °C with constant shaking. Equal volumes of extracted RNA were used as input for RT-qPCR. Data were analysed as described in Gagnon et al. (26) with minor modifications. Briefly, data were normalised relative to GAPDH levels in each cellular compartment and the fold change was calculated compared to the whole cell lysate. As miR-27 RT-qPCR was performed with the miRCURY kit that is not designed for mRNA quantification, miR-27 levels were only normalised to the whole cell lysate.

### Western blot

Cells were lysed in CLEAR-CLIP lysis buffer or processed as described for nuclear fractionation. Same volumes, equivalent to 1×10^5^ cells were mixed with 4x NuPAGE LDS sample buffer (Invitrogen) and run on a 4-12 % NuPAGE Bis-Tris protein gel in MOPS SDS running buffer (Invitrogen). Proteins were then transferred onto a PVDF membrane (Immobilon-FL, Merck) at 100 V for 105 min. Membranes were blocked with 3 % milk powder in TBS + 0.1 % Tween-20 (TBS-T) and then incubated with the primary antibody overnight at 4 °C. Primary antibodies used in this study: AGO2 (1:1500, clone C34C6, Cell signalling, 2897), α-Tubulin (1:2000, Cusabio, CSB-PA004344), β-Actin (1:1000, Cell signalling, 4967), Calreticulin (1:2000, Cell signalling, 2891), CEACAM1 (1:20,000, Abcam, ab235598), Histone H3 (1:2000, Cell signalling, 4499), ZFAND5 (1:1000, Sigma, HPA018129). The next day the membrane was washed with TBS-T and incubated with the secondary antibody (Goat anti-Rabbit IgG (H+L) Secondary Antibody, DyLight 800, Thermo Scientific) diluted 1:20,000 in TBS-T containing 3 % milk or 5 % BSA for 1 h. Membranes were washed again before protein bands were visualised using the Odyssey CLx scanner (Li-cor). For quantification of protein levels, the bands were quantified using Fiji ImageJ (27) as outlined in Stael et al. Section 3.5 (28).

### Northern blot

To detect RNAs of interest by Northern blot, 1 µM of the probes (Table S9) were 5’ end labelled with [γ^32^P] ATP (6000 Ci/mmol, Perkin Elmer) using the T4 polynucleotide kinase (10 U/μL, Thermo Scientific) in a total volume of 20 µL and subsequently cleaned up with Microspin G-25 columns (GE Healthcare) according to the manufacturer’s instructions. Equal amounts of extracted RNA (3.5 µg) were mixed 1:1 with Gel Loading Buffer II (Invitrogen) and loaded on a 15 % polyacrylamide/8 M Urea gel. RNA Decade Marker (Thermo Scientific) was prepared according to the kit instructions, diluted in Gel Loading Buffer II, and run along the samples. The gel was run in 0.5× TBE at 120 V for 2 h. Samples were then transferred onto a Hybond-N membrane (Amersham Cat. RPN203N, 1 h at 80 V in 0.5× TBE) and chemically crosslinked at 55 °C for 2 h using a solution of 0.16 M EDC (N-(3-Dimethylaminopropyl)-N′-ethylcarbodiimide hydrochloride, Merck) with 0.13 M 1-methylimidazole in H2O (pH 8).(29) The membrane was washed once with 2× SSC (20× SSC, Fisher Scientific) and then pre-hybridised in 4 mL of Perfecthyb Plus hybridisation buffer (Sigma-Aldrich). 5’ end labelled probes were heated to 90 °C for 30 s and hybridised to the membrane over night at 42 °C with rotation. The next day the membrane was washed once with the following pre-heated buffers: Wash buffer 1 (2× SSC, 0.1 % SDS), wash buffer 2 (1× SSC, 0.1 % SDS) and wash buffer 3 (0.1× SSC, 0.1 % SDS). To visualise the RNA, the membrane was incubated on a Storage Phosphor screen for 24 h before it was imaged on a Typhoon Scanner (GE Healthcare).

### Bioinformatic analysis

### Small RNA sequencing analysis

Reads were first quality checked using FastQC (http://www.bioinformatics.babraham.ac.uk/projects/fastqc/) and low quality 3’ bases (quality below 20) were removed. The 3’ small RNA adapter TGGAATTCTCGGGTGCCA was trimmed using Cutadapt (version 1.15) (30) and only reads containing the adapter and ≥ 18 nt were retained. miRNAs were identified using QuickMIRSeq (31) with standard settings. The QuickMIRSeq output file with miRNA counts was then uploaded on Degust web tool (Version 4.1.3, Powell, 2015, DOI: 10.5281/zenodo.3258933) for differential expression analysis using the Voom/Limma method with no filters or cut-offs. Differential expression in RSV vs Mock samples was visualised using ggplot2 R package and only changes with an FDR < 0.05 were considered significant.

## CLEAR-CLIP

Initial processing (quality control and trimming) was performed as described above. For paired-end libraries only the forward read was used in the analysis, since it contains sufficient sequence for identifying chimeric reads. All reads were mapped to mature Human miRNAs (miRbase V22) with bowtie2 using local sensitive parameters (--sensitive-local), as this allows for partial alignment of potentially chimeric reads to the miRNA database, as well as identifying non-chimeric miRNA reads (32). If the first part mapped to a miRNA and the remaining part was ≥ 18 nt and mapped to the human genome, this read was classified as miRNA-target-chimera. For the initial round of sequencing (NovaSeq 6000, 100-base, single end – except for 06_Mock (CLEAR_200_UP), see Table S1) we mapped the target part of the read to all human protein coding transcripts (GRCh38.p7, Gencode) using bowtie2. For the final dataset (NovaSeq 6000, 150-base, paired-end, as well as 06_Mock (CLEAR_200_UP), see Table S1), the target part was aligned to the human genome (GRCh38.p7, Gencode) with bowtie2, then the aligned portion was extracted and removed if it matches a low complexity sequence as determined by dustmasker. Those target read portions that were kept were then re-mapped to the human genome using ShortStack (33) as follows: allowing 1 nt mismatches, requiring clustered-reads to have a minimum expression level of 0.5 reads per million (--mincov 0.5 rpm), all potential alignment sites were considered for multimapping reads (--bowtie_m ‘all’) with no upper limit on number of locations (--ranmax ‘none’). If a read mapped to multiple locations, nearby uniquely mapped reads are used to guide the chosen location (--mmap u) and when no uniquely mapped reads are available the location is chosen randomly. Once reads were distributed, overlapping reads were merged into clusters with bedtools v2.30.0 using the merge functionality with the parameters –s and –o ‘distinct’. Genome features were obtained from Ensembl regulation release 100, and included in the analysis were 3’ untranslated regions (UTRs), coding sequences (CDS), 5’ UTRs, introns, regulatory regions (CTCF binding sites, transcription factor (TF) binding sites, enhancer regions, open chromatin regions, promoter-flanking regions and promoter), unprocessed and processed pseudogenes, tRNAs, lncRNAs, rRNAs and miRNAs. Overlaps between annotations and the chimeric cluster regions were found with bedtools (34). We then applied a set of filters to identify high confidence chimeras among these interactions. Firstly, target sites need to be present in ≥ 2 samples of the same condition (RSV-infected or mock-infected). Secondly, as the requirement of only a 1 nt overlap can lead to exceedingly long target sites with low overall overlap, the target site length should be shorter than 150 nt, and thirdly the target site must contain a miRNA seed site with no mismatches. The code used for this part of the analysis is available on Zenodo (https://doi.org/10.5281/zenodo.7924850). Chimeric miRNA-RSV interactions were identified as described above, except the target part of the read was required to not map to the human genome and map to the RSV genome (KT992094). miRNA-target binding patterns were predicted using the IntaRNA web tool (35). Raw data as well as a processed table with all miRNA-host target interactions have been deposited in GEO under accession GSE232686. Re-analysis of samples from the Moore *et al.* study (16) was performed by first removing degenerate adapters to make those samples comparable to our data. We generated separate bedgraph files for each miRNA detected in the CLEAR-CLIP data. Two different sets of tracks were made: the first one included all interactions and the second one only high-confidence interactions as defined above. The tracks were made using R 4.2.2 from alignment bam files using either the human or RSV genome as reference. The packages used to generate the tracks were: Rsamtools 2.14.0 (DOI:10.18129/B9.bioc.Rsamtools) for importing bam files; dplyr 1.1.2 to apply the high confidence filter; GenomicRanges 1.50.2 (36) to find overlaps between the bam files and high confidence target clusters as well as rtracklayer 1.58.0 (37) to export bedGraph files. For human tracks bedgraph files were transformed to TDF format with igvtools 2.5.3 (38).

### Total RNA-seq

Read quality and adapter trimming was performed with trimmomatic 0.39 (39) using the TruSeq3 PE adapters. Paired-end reads were aligned to the human genome (GRCh38.p7, Gencode) with hisat2 2.2.1 (40). Visualization tracks were made within R with the packages: GenomicAlignments 1.28.0 (36), GenomicRanges 1.44.0 (36), Rsamtools 2.8.0 (DOI:10.18129/B9.bioc.Rsamtools), rtracklayer 1.52.0 (37). Each bam file was read into R with the readGAlignmentPairs function without reading secondary alignments; for the total RNA-seq tracks of Chr 19 only unique mapping reads were considered. The strand mode for the GAlignmentPairs object was set to 2. Reads were split by plus or negative strand and coverage was calculated separately; the coverage of the negative strand was multiplied by –1 to show reads below the line. Both coverages were then joined in a single object and saved as bedGraph files, which were subsequently transformed with igvtools (38) to TDF format.

A custom GFF3 file was built which included gene annotations available through Gencode and regulatory regions provided by Ensembl regulation (release 100); both sources of annotations were merged in R using rtracklayer 1.52.0. All regulatory regions that overlapped any Gencode gene annotations were removed, and the feature type was set to “exon” to allow regulatory regions to be quantified in downstream processes. The number of reads mapping to genomic features were quantified with the featureCounts function implemented in Rsubread (version 2.6.4) R package (41) using the following non-default parameters: isPairedEnd = TRUE, isGTFAnnotationFile = TRUE, fraction = TRUE, largestOverlap = TRUE, strandSpecific = 2.

The upset plot for multiple overlapping annotations was made with the ComplexUpset 1.3.5 R package. Degust web tool (Version 4.1.3, Powell, 2015, DOI: 10.5281/zenodo.3258933) was used for differential expression analysis using the Voom/Limma method, with min gene read count of 4 and min gene CPM of 1 in at least 1 sample. Genes were considered differentially expressed with an FDR < 0.05. CDF plots were made with ggpubr R package. Target genes with more than one binding site were collapsed to ensure each gene was only present once per category.

### Statistical tests

Details regarding the statistical tests used and the represented data can be found in the figure legends, or, for the differential expression analysis in the “Small RNA sequencing”/ “Total RNA-seq” section of the Bioinformatic analysis. Statistical tests were considered significant when P values were less than 0.05. Statistical analyses were performed via GraphPad Prism (Prism Software) or, for the Cumulative Distribution Function (CDF) plots in RStudio using the wilcox.test function with alternative = “two.sided” except for mimic and inhibitor plots, for which alternative = “less” (“greater” for inhibitor data) was used. For the statistic comparison of HC targets vs all targets, HC targets were removed from the “all targets” data set.

## RESULTS

### CLEAR-CLIP protocol optimisation in A549 cells

To generate a high quality CLEAR-CLIP dataset in A549 cells, we tested and modified several parameters compared to the protocols reported previously (4,16,20). The first step of the protocol involves UV crosslinking to generate covalently linked protein-RNA complexes, where different doses have been used across different cellular contexts (14,16,20,42,43). We found a suitable dose in A549 cells to be 200 mJ/cm^2^. In contrast, no crosslinking yielded fewer chimeric reads and was enriched for miRNA reads, as expected from previous literature (16). Higher doses (400 mJ/cm^2^) were associated with protein loss and fewer miRNA-target chimeras (Fig S1A-C, Table S1). Irradiated cells were immediately lysed and subsequently treated with RNases and the intermolecular ligation step was carried out with minor modifications (see Materials and Methods). A 3’ adapter with an infrared dye allowed for visualisation of the RNA on SDS-PAGE (Fig S1D) and excision of the AGO-RNA complexes was carried out from the gel without the need for transfer to a membrane. We analysed two different molecular weight regions and found the high molecular weight AGO2 enriched for miRNA-target interactions (115-185 kDa, Fig S1C), consistent with previous reports (16,44).

Replicate datasets were generated with A549 cells (1×10^8^) either infected with RSV at a multiplicity of infection (MOI) of 0.1 or mock-infected for 48 h (for summary of protocol see Fig 1A). Chimeric reads were identified based on the requirement that the first part of the read maps to a mature miRNA. The remaining part of the read is then used to identify targets with a bioinformatic pipeline that maps reads to the human genome using Shortstack (33) and clusters mapped reads into target sites (Fig 1B, S1H). The percentage of chimeric reads with the human genome ranged from 3-6 %, on par with previous reports in other cell types (Fig 1C). By mapping to the genome instead of the transcriptome our analysis can in principle identify target sites beyond mRNAs (CDS/UTRs). A resulting challenge of this approach is the large number of overlapping genome annotations (here classified as “Multiple”, Fig S1E). Most analyses apply an annotation hierarchy to solve this problem. As target sites in 3’UTRs have been shown to be the most functional miRNA targets, we too classify “multiple” target sites that overlap 3’UTR as 3’UTRs for downstream analysis (> 50 % of the “multiple” target sites overlapped 3’UTRs (Fig 1D)). However, the remaining target sites with overlapping annotations were kept in the “multiple” category and not further classified (Fig 1E, Fig S1F). The initial analysis of seven datasets (n=3 in Mock and n=4 in RSV-infected) yielded a total of 389,168 and 220,938 unique target sites across Mock and RSV-infected samples, respectively. As a gauge for potential non-specific chimeras that could occur during sample preparation, we note that 10-18 % of the target sites were mapped to ribosomal RNAs (rRNAs, Fig 1E, S1G); this is comparable to a similar percentage of rRNA chimeric reads in other CLEAR-CLIP reports: 12-17 % in Moore et al. (16) based on re-analysis of their data with our pipeline (Fig S1G).

**Figure 1.**
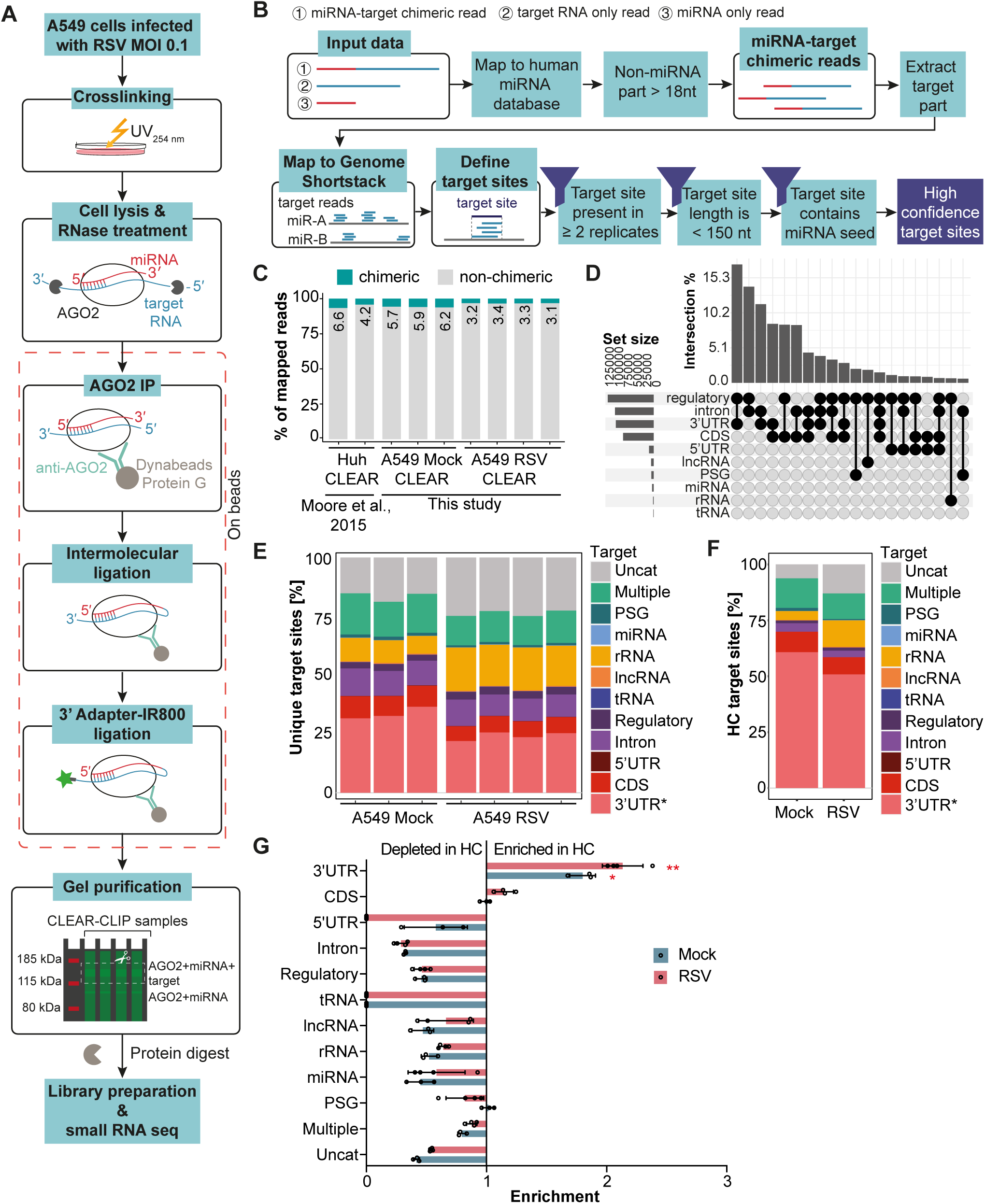
miRNA target sites in RSV-infected A549 discovered by CLEAR-CLIP. (**A**) Overview of the CLEAR-CLIP protocol. (**B**) Schematic of bioinformatics pipeline: chimeric reads were defined as the first part of the read mapping to a mature miRNA with the remaining part ≥ 18 nt. Target sites were extracted and mapped to the human genome and subsequently clustered into target sites per miRNA. To remove background of non-specific interactions, target sites were filtered for reproducibility, length < 150 nt, and presence of the miRNA seed. (**C**) Percentage of chimeric vs non-chimeric reads mapping to the human genome in our samples (n = 3 for Mock and n = 4 for RSV) compared to the Moore et al. study (16). (**D**) Upset plot of target sites with multiple annotations. Annotations are regulatory regions (regulatory), introns, 3’ untranslated regions (3’UTR), coding sequence (CDS), 5’UTR, long non-coding RNAs (lncRNA), pseudogenes (PSG), miRNAs, ribosomal RNAs (rRNA), and transfer RNAs (tRNA). Intersections were filtered for top 10. (**E**) Percentage of unique target sites across different RNA categorises as described in (D) as well as multiple annotations (Multiple) and uncategorised (Uncat). All target sites overlapping 3’UTRs were classified as 3’UTRs in this graph (thus removing those from Multiple category). (**F**) Same as in (E) but showing the percentage of unique high confidence (HC) target sites. (**G**) enrichment of HC target sites compared to non-filtered target sites. Red asterisks indicate significant enrichment (One Brown-Forsythe and Welch ANOVA with Dunnett’s T3 multiple comparisons test performed separately for Mock and RSV samples, * P ≤ 0.05, ** P ≤ 0.01).

### Filtering chimeric reads for reproducibility, cluster length and seed sites enriches for miRNA-target interactions in 3’UTRs

Previous studies have shown that AGO scans for targets in a stepwise mechanism, searching first for complementarity with nucleotides 2-4 of the miRNA before extending complementarity to the nucleotides 5-8 (45). To reduce the likelihood of non-specific and non-functional targets in our dataset, we required that target sites were reproducible in ≥ 2 Mock or ≥ 2 RSV samples, and that the length of the cluster of target reads was < 150 nt (for reference clusters within ribosomal RNAs have a mean of >150 nt whereas those within 3’UTRs are <100 nt; Fig S1H, I). We then applied a further filter to require that target interactions contain canonical seed-based pairing (not allowing mismatches or bulges but allowing 6mer-offset as defined in Bartel (46)) since miRNAs are generally thought to require the interaction with the seed site for efficient inhibition of target translation (46). Our filtering steps of reproducibility, length and seed site resulted in a subset of 7,539 and 2,898 high confidence (HC) target sites in Mock and RSV-infected samples respectively, with over 50 % of the target sites mapping to 3’UTRs (Fig 1F). The HC target sites were significantly enriched for 3’UTRs (compared to the unfiltered chimeric target list) with other target types being depleted, including rRNAs (Fig 1G).

### miRNAs miR-26 and miR-27 interact with specific sites in the RSV genome but are not degraded upon infection

To examine whether RSV interacts directly with host miRNAs, we mapped non-chimeric (AGO binding) and chimeric reads from the CLEAR-CLIP datasets to the RSV genome. We find that chimeric reads with RSV make up on average 0.09 % of the total mappable reads in infected cells (Fig S2A) compared to 3.3% for chimeric reads with host genes (Fig 1C). Chimeras and AGO binding sites were found across most RSV genes but were by comparison underrepresented in NS1 and L genes (Fig 2A, B). This appears to show selectivity as transcripts for NS1 and L genes are expressed at comparable levels based on RNA sequencing (Fig 2A, top panel). However, we note that a large portion (89.1%) of RSV-miRNA chimeras are neither reproducible nor seed-mediated, and the majority are comprised of interactions with miR-21 which ranks first in abundance in these cells based on small RNA sequencing (Fig 2B-D, Table S2). To reduce scope for non-specific interactions, we further required that a miRNA interact in a reproducible, seed-based manner. Figure 2B shows the interactions for the top 10 miRNAs with the most HC chimeric read counts. We found that miR-26 and miR-27 mapped specifically to individual sites in the M gene (Fig 2B, C). miR-26 has the most interactions with the RSV genome that are reproducible and seed-based (Fig 2C, Table S2). We note that neither miR-26 family member is among the most abundant miRNAs in these cells (miR-26a ranks 13^th^ and miR-26b 28^th^) based on small RNA sequencing (Fig 2D). While miR-27 is more abundant in the cell (miR-27b ranks 4^th^ and miR-27a ranks 45^th^ based on small RNA sequencing, Fig 2D) the pattern of binding is highly specific in comparison to miR-21 (Fig. 2C).

**Figure 2.**
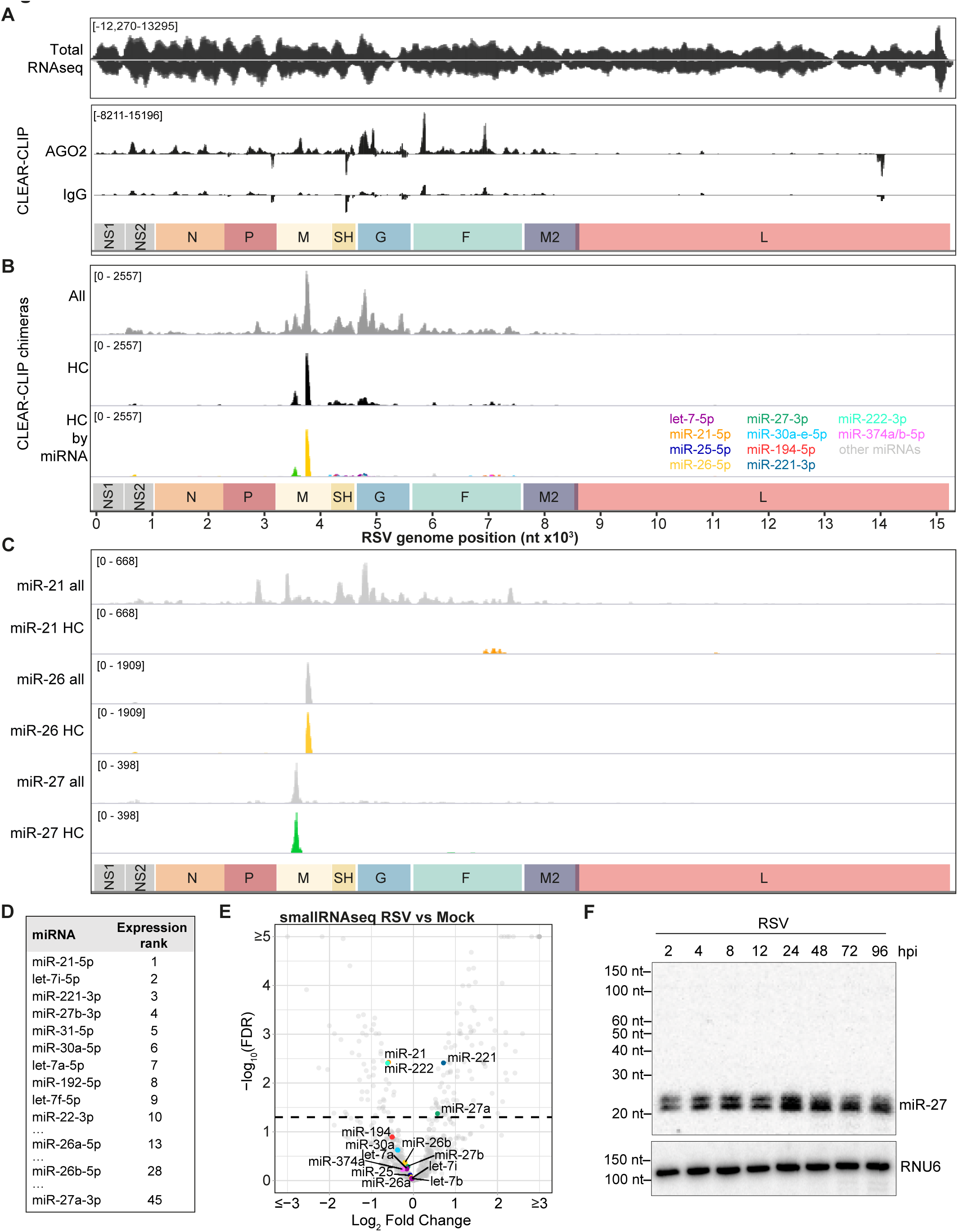
miR-26 and miR-27 interact with specific sites in the RSV genome but are not degraded upon infection. (**A**) Genome browser view of the RSV genome (KT992094) showing total RNA-seq read coverage (top panel), as well as AGO2 and IgG binding (determined from non-chimeric reads mapping to RSV, lower panel). (**B**) Same as in (A) but showing all miRNA chimeras (All, top), reproducible, seed-based miRNA chimeras (HC, middle), and reproducible, seed-based binding sites for the top 10 miRNAs interacting with RSV genome (determined from HC read counts, HC by miRNA, bottom). (**C**) Same as in (A) but showing all interactions with RSV genome compared to reproducible, seed-based chimeras (HC) for miR-21, miR-26, and miR-27. (**D**) Table showing the top 10 most highly expressed miRNAs in A549 cells as well as all miR-26 and miR-27 family members, ranked by mean expression in RSV samples determined from smallRNA-seq. (**E**) Volcano plot of smallRNA-seq data showing the log_2_ fold change in miRNA expression in RSV-infected samples compared to Mock-infected samples. Dots representing miRNAs highlighted in (C) are coloured and labelled. Dashed line indicates FDR 0.05. For illustration purposes axes were limited to log_2_ fold change of –3/3 and –log_10_(FDR) of 5. Points outside of these limits were assigned the maximum values. (**F**) Northern blot showing miR-27 length over time course of RSV infection compared to loading control RNU6.

Previous studies have shown that interactions with viral RNA can lead to miRNA degradation, in particular in the case of two different herpesviruses targeting miR-27 (5–8) where the viral RNAs exhibit extensive pairing with the 3’ end of the miRNA (47). To determine whether miR-27 or miR-26 were degraded during infection, we examined miRNA differential expression by small RNA-seq. The expression of miR-26 and miR-27b does not significantly change in RSV infection, while miR-27a is slightly upregulated at 48 hpi (hours post infection, Fig 2E). In addition, the interactions of both miRNAs with RSV do not involve 3’ end pairing (Fig S2B, C) which is required for miR-27 degradation by herpesviruses (48). However, we note that miR-27 is shortened over the time course of RSV infection from 21 nt to 20 nt (Fig 2E, S2D) – which is not observed for miR-26 (Fig S2E). The effect that this shortening would have on target interaction is not clear, but it has been proposed that 3’ end variations affect AGO2 binding as well as miRNA-mRNA interactions (49).

### miR-26 and miR-27 target distinct host genes and networks regulating cell cycle, metabolism, and the antiviral immune response

To determine whether the interactions of miR-26 and miR-27 with RSV might sequester these miRNAs from their host targets, we first compared the read counts of all HC targets in uninfected and infected cells. We note that there is a general shift towards a reduction in HC chimeric reads upon infection (black line, Fig 3A, B). This is consistent with less overall chimeras being sequenced from infected cells and slightly more background (rRNA) reads (Fig 1E, F, Table S1). Yet miR-27 HC chimeric reads are significantly reduced upon infection compared to all others (Fig 3A; two-tailed Wilcoxon rank sum test p-value=3.6×10^-16^). Strikingly, we find the opposite result with miR-26 HC chimeric reads: there are significantly more in infected versus uninfected cells (Fig 3B; two-tailed Wilcoxon rank sum test p-value=3.5×10^-5^).

**Figure 3.**
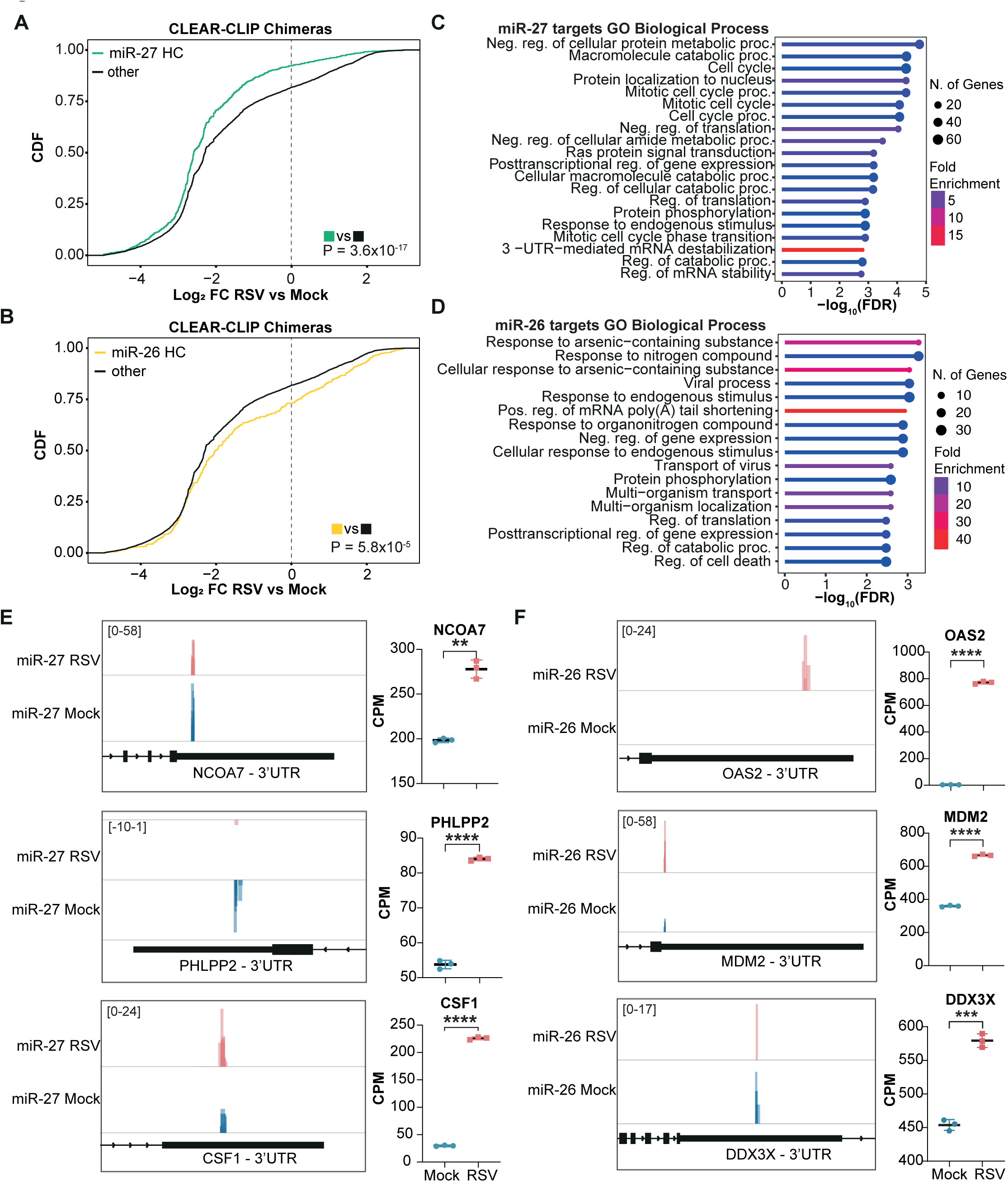
miR-26 and miR-27 target distinct host genes and networks associated with cell cycle, metabolism and antiviral immune response that are upregulated upon infection. (**A, B**) Cumulative distribution function (CDF) plot of CLEAR-CLIP host target interactions comparing log_2_ fold change (Log_2_ FC) in RSV vs Mock samples of miR-27 HC interactions (**A**) or miR-26 HC interactions (**B**) compared to HC interactions of all other miRNAs. Two-tailed Wilcoxon p-values are shown for the comparisons. (**C, D**) Pathway analysis of miR-27 (**C**) or miR-26 (**D**) HC 3’UTR target genes using ShinyGO V0.77 (50) showing the top significant pathways (FDR < 0.05, pathway size: min. 2, max. 2000) for GO term Biological process. (**E**) Genome browser view showing HC interaction sites of miR-27 in Mock and RSV-infected cells for NCOA7, PHLPP2, and CSF1 as well as gene expression levels in counts per million (CPM, right) determined from total RNA-seq in Mock and RSV-infected cells, shown as mean ± SD. Significance of changes in gene expression were determined using unpaired t-tests with Welch’s correction (** P ≤ 0.01, *** P ≤ 0.001, **** P ≤ 0.0001). (**F**) Same as for (E) but showing HC interaction sites of miR-26 with OAS2, MDM2, and DDX3X.

To understand the nature of the infection-induced differences in target engagement we examined the 395 HC 3’UTR targets of miR-27 and 165 HC 3’UTR targets of miR-26 that we identified in A549s (Table S3). We note these are reproducible, seed-based interactions where 22 % (miR-27) and 39 % (miR-26) of those we identify here overlap validated human targets on miRTarBase (50) that come from diverse cellular contexts (Fig S2F, G). Strikingly, GO term analysis using ShinyGO (51) showed no overlap in the top 10 most significantly enriched pathways for miR-26 and miR-27 HC targets (Fig 3C, D), suggesting these two miRNAs may have distinct functions in the cell and supporting the specificity of the target interactions that we identify here. The top most enriched pathways for miR-27 HC targets were “metabolic processes” and “cell cycle” (Fig 3C, Table S5), while for miR-26, we found processes associated with viral infection (“viral process” and “transport of virus”, Fig 3D, Table S6). Several miR-27 HC targets fit a model consistent with de-repression (less chimeras in infected cells and higher gene expression upon infection), for example *NCOA7* (Nuclear receptor coactivator 7) that is associated with metabolism and *PHLPP2* (PH Domain And Leucine Rich Repeat Protein Phosphatase 2) that regulates cell cycle (Figure 3E, Table S4). However not all HC miR-27 targets show this pattern; an example is *CSF1* (Colony Stimulating Factor 1, Fig 3E, left) which is involved in intercellular signalling and macrophage activation (52). We identify more miR-27 HC chimeras with *CSF1* in infected cells compared to uninfected cells, which could be due to transcriptional up-regulation of this gene upon infection (Fig 3E, right, Table S4).

As detailed above, many individual miR-27 targets are up-regulated upon infection, but globally this is not a significant trend compared to other genes that are not miR-27 targets (Fig S2H). However, there is a trend towards upregulation for miR-26 targets (Fig S2I). This is consistent with the pathway analysis which shows miR-26 HC targets are enriched in pathways including “viral processes”. Examples include *OAS2* (2’-5’-Oligoadenylate Synthetase 2) which is involved in the innate immune response to viral infection (53) and only expressed in infected cells and *MDM2* (Mouse double minute 2 homolog) (Fig 3F), a known negative regulator of p53 that is linked to antiviral immunity (54,55). *DDX3X* (DEAD-box helicase 3 X-linked) is a miR-26 target that upon infection increases in expression but has fewer HC chimeras (Fig 3F). *DDX3X* is particularly interesting because it contains two adjacent miR-26 sites in the 3’UTR (Fig S2J) and this host protein is known to be required in numerous viral life cycles (56). Although binding of miR-26 to RSV could contribute to the de-repression of *DDX3X* upon infection we note that the other mentioned miR-26 targets do not fit this model: many targets have increased HC chimeric reads with miR-26. Collectively, the analysis of HC chimeras solidifies miR-26 and miR-27 as regulators of key genes important in viral infection including those associated with cell cycle regulation, metabolism, and antiviral immunity. Our data do not support a model where binding of the miRNAs to RSV leads to a global de-repression of all host miRNA targets and highlights the fact that different targets could have different susceptibilities to inhibition, and some targets will not even be present prior to infection.

### miR-27 HC targets that are most sensitive to miR-27 inhibition are involved in cell cycle regulation and metabolism

To explore further the sensitivity of host targets to miRNA inhibition, we examined how the HC 3’UTR targets are impacted when cells are treated with miR-27 inhibitors and in comparison to treatment with miR-27 mimics. Total RNA-seq showed that overexpression of miR-27 resulted in downregulation of miR-27 HC 3’UTR targets compared to controls (Fig 4A) while miR-27 inhibition resulted in a significant upregulation of miR-27 HC targets (Fig 4B). In all the conditions the miR-27 HC 3’UTR targets (395 genes) were more strongly and significantly regulated compared to all of the miR-27 3’UTR targets (4415 genes), highlighting the value of our filtering approach to identify functional targets. Overall, we found 181 miR-27 HC targets to be significantly downregulated by the mimic and 37 HC targets to be significantly upregulated by the inhibitor (FDR < 0.05, Table S7).

**Figure 4.**
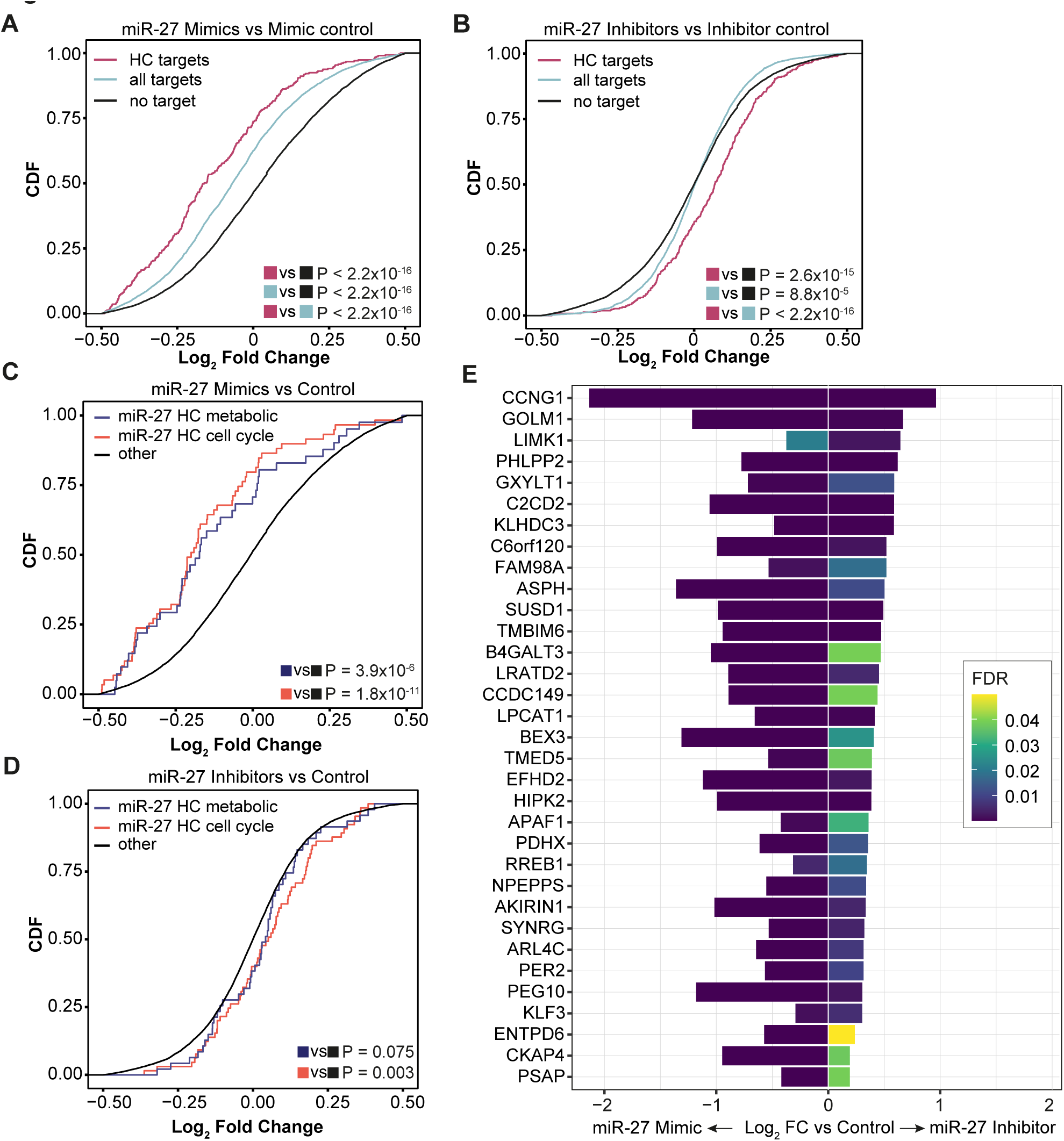
miR-27 inhibition regulates genes associated with cell cycle progression and metabolism. (**A-B**) CDF plots of RNA-seq data comparing the log_2_ fold change of miR-27 HC 3’UTR target genes (red) vs all miR-27 3’UTR target genes (light blue) and no target genes (black) between A549 cells transfected with 10 nM miR-27 mimics or control mimic (**A**), and miR-27 inhibitors or control inhibitor (**B**). One-tailed Wilcoxon p-values are shown for the different comparisons. (**C-D**) CDF plots of RNA-seq data comparing the log_2_ fold change of miR-27 HC 3’UTR target genes associated with either metabolic processes (metabolic) or cell cycle against all other genes in A549 cells transfected with miR-27 mimics vs control mimic (**C**), or miR-27 inhibitors vs control inhibitor (**D**). One-tailed Wilcoxon p-values are shown for the different comparisons. (**E**) Bidirectional bar plot showing the log_2_ fold change of miR-27 HC 3’UTR target genes significantly (FDR < 0.05) downregulated upon miR-27 mimic transfection compared to control AND significantly upregulated upon miR-27 inhibitor transfection compared to control. Log_2_ fold change < 0 shows change with mimic, log_2_ fold change > 0 shows change with inhibitor vs respective controls. Genes are ranked by log_2_ fold change of miR-27 inhibitor vs control. Colour indicates FDR.

We confirmed that the miR-27 HC targets associated with cell cycle and metabolism pathways (from Table S5) are functionally regulated by both inhibitor and mimic (Fig 4C, D). Analysis of the top up-regulated targets by the miR-27 inhibitor further highlights cell cycle, metabolism, and antiviral immunity targets as susceptible to miR-27 inhibition. The top four targets that are significantly upregulated are: CCNG1 (Cyclin G1) and PHLPP2 which have both been shown to induce cell cycle arrest (57–61), GOLM1 (Golgi membrane protein 1, also known as GP73) which is a validated miR-27 target that is emerging as a key regulator of IL-6 also linked to innate immunity and RSV infection (62–66) and LIMK1 (LIM domain kinase 1, Fig 4E), a serine/threonine-protein kinase involved in remodelling of the cytoskeleton. LIMK1 has been shown to be an important host factor for many viral life cycles, including HIV, HSV-1, and Influenza virus (67–71) and LIMK1 inhibitors are currently explored as broad-spectrum antiviral drugs (72). These data suggest that the miR-27 HC targets most susceptible to miRNA inhibition in lung epithelial cells are highly relevant to viral infection.

### Pseudogenes and lncRNAs are HC miRNA targets and nuclear localisation of AGO2:miR-27 suggest potential roles in transcriptional gene regulation

The above analyses solidify and expand the role of miR-26 and miR-27 in regulating 3’UTR targets associated with multiple pathways that impact viral infection. Yet these targets still may not reveal the whole story of how the miRNAs could impact infection. Recently, there has been increasing evidence that miRNA functional mechanisms are not limited to 3’UTR targeting and it has been shown that AGO:miRNAs can also regulate transcription through interaction with promoter-associated lncRNAs, leading to either activation or inhibition of gene expression in the nucleus (73–77). We therefore wished to take advantage of our chimeric data and HC filters to determine whether we find evidence for targets mapping to regulatory regions. The caveat/challenge of our approach (mapping to the genome rather than the transcriptome) is that a given region of the genome can have more than one bonafide annotation, e.g. some 3’UTRs can overlap regulatory regions (Fig 1D). We therefore removed all targets sites that map to 3’UTR and CDS prior to examining the remaining HC target sites. As shown in Figure 5A, regulatory regions that overlap pseudogenes and lncRNAs were found to be the top two most enriched categories in HC target sites (Fig 5A). In total, we found 153 unique HC target sites located in RNAs from regulatory regions, overlapping either PSGs or lncRNAs, 10 of these are miR-27 chimeras and 5 are miR-26 chimeras (Table S8). Here we highlight one example for miR-27: a highly conserved promoter flanking region (reannotated as enhancer in Ensembl release 109) that overlaps *zinc finger, AN1-type domain 5* (*ZFAND5*) processed PSG on chromosome 19. Although it is challenging to distinguish reads that map to the PSG compared to the parental gene, we note that in this case unique mapping reads from RNA-seq confirmed that a lncRNA is expressed from the PSG locus (Fig 5B, S3A) and this finding is supported by a previous study analysing data from the Genotype-Tissue Expression (GTEx) project (78). The highly conserved miR-27 binding site is located in the CDS of the PSG as well as the parental gene (the PSG does not contain introns or UTRs, Fig S3A) and we find a let-7 site adjacent to the miR-27 site (Fig 5B). We validate that the lncRNA derived from the PSG is highly enriched in the nuclear fraction (to a similar extent as the nuclear control lncRNA Malat1) and in comparison to the parental ZFAND5 (Fig 5C). We also find that miR-27 is present in both nuclear and cytoplasmic fractions, although we detect a small proportion of the cytoplasmic control RPL30 in the nuclear fraction (suggesting that nuclear proportion of the RNAs might be slightly overestimated in our analysis). Using western blot analysis, we show that AGO2 is present in the nuclear fraction (Fig 5D, E) in both naïve or infected cells and that a small proportion of AGO2 is associated with the chromatin containing fraction (Nuc insoluble, Fig 5F). These data are consistent with previous work showing nuclear localisation of AGO2 in lung epithelial cells (26) and support the possibility that miRNAs and AGO2 could be involved in transcriptional regulation in these cells. Finally, to test potential functionality of the ZFAND5 PSG lncRNA we used GapmeRs to knock down the lncRNA (Fig 5G) and demonstrate an increase in the protein levels of CEACAM1 (Fig 5H, I), a gene that is located proximal to the PSG lncRNA on chromosome 19 (Fig S3B). Although the GapmeR also reduces the parental ZFAND5 expression (Fig S3C), we show that ZFAND5 itself does not regulate CEACAM1 since siRNA knockdown of *ZFAND5* did not affect CEACAM1 protein levels (or the levels of the PSG lncRNA, Fig 5H, I, S3D). This work is consistent with a model where the lncRNA derived from the PSG could repress transcription of nearby genes. Collectively, our analyses expand the type of miR-27 interactions that exist in RSV-infected cells and suggest that target sites beyond those in the 3’UTR may merit further attention when assessing the contributions of miRNAs to gene regulation during viral infections.

**Figure 5.**
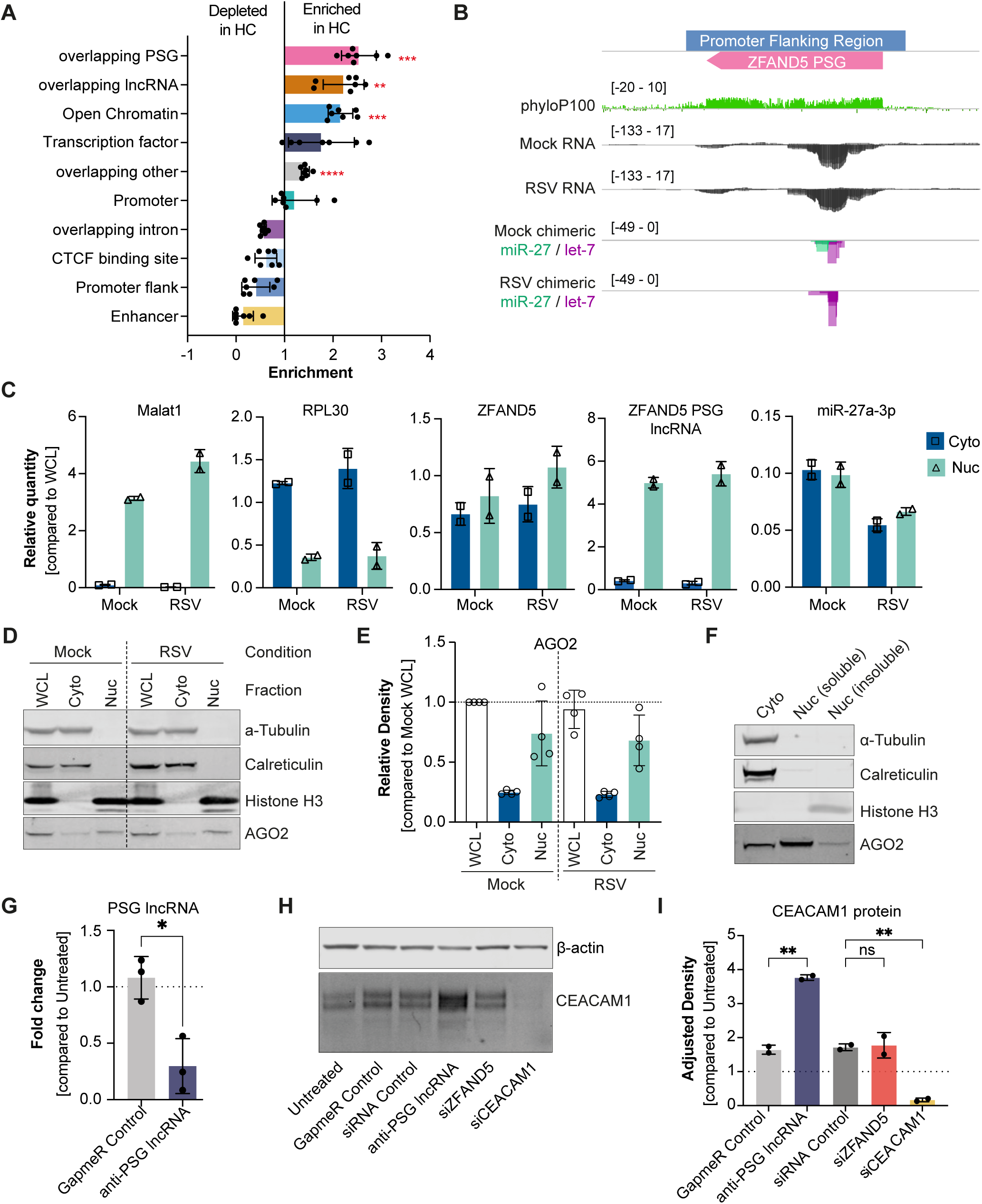
Pseudogenes and lncRNAs are HC miRNA targets and nuclear localisation of AGO2-miR-27 suggests potential roles in transcriptional gene regulation. (**A**) Enrichment of HC target sites in regulatory regions compared to non-filtered target sites. Target sites were filtered to remove sites overlapping 3’UTRs and CDS. Shown are the mean ± SD for three Mock and four RSV CLEAR-CLIP replicates combined. Red asterisks indicate significant enrichment; One Brown-Forsythe and Welch ANOVA with Dunnett’s T3 multiple comparisons test. Due to low number of target sites in some of the categories, Mock and RSV samples were analysed together for more statistical power, * P ≤ 0.05, ** P ≤ 0.01, *** P ≤ 0.001, **** P ≤ 0.0001. “Overlapping” indicates regulatory regions overlapping other genome annotations (pseudogenes (PSG), lncRNAs, introns, and other). (**B**) Genome browser view of the genomic location of top miR-27 regulatory target overlapping a PSG, showing the conservation using PhyloP100, features of the genomic locus, total RNA-seq coverage in Mock and RSV-infected samples (n = 3), as well as miRNA target sites coverage in Mock and RSV-infected CLEAR-CLIP samples. (**C**) RT-qPCR results for Malat1, RPL30, ZFAND5, ZFAND5 PSG lncRNA, and miR-27a (left to right) levels in cytoplasmic (cyto) and nuclear (nuc) fractions in Mock and RSV-infected cells. Data is shown as mean ± SD for two biological replicates. (**D**) Western blot analysis of AGO2 protein levels in subcellular fractions showing whole cell lysate (WCL), cytoplasmic (cyto), and nuclear (nuc) fractions for Mock and RSV-infected cells. α-Tubulin and Calreticulin were used as cytoplasmic markers and Histone H3 as nuclear marker. (**E**) Quantification of AGO2 protein levels in (D). Data is shown as mean ± SD for four biological replicates and normalised to WCL Mock. (**F**) Western blot analysis of cellular fractionation of naïve A549 showing cytoplasmic fraction as well as the soluble and insoluble nuclear fractions. α-Tubulin and Calreticulin were used as cytoplasmic markers and Histone H3 as nuclear marker associated with chromatin. (**G**) RT-qPCR results of PSG lncRNA expression after treatment with 5 nM GapmeR targeting PSG lncRNA (anti-PSG lncRNA) or control. Shown are three biological replicates with mean ± SD. Significance was tested with unpaired two-tailed t-test (* P ≤ 0.05). (**H**) Western blot analysis of CEACAM1 after treatment with anti-PSG lncRNA GapmeR, siRNAs against ZFAND5 (siZFAND5) and CEACAM1 (siCEACAM1) and controls compared to β-Actin loading control. (**I**) Quantification of CEACAM1 protein levels from (H) corrected for β-Actin and normalised to Untreated control. Shown are two biological replicates with mean ± SD. Significance was tested with One Brown-Forsythe and Welch ANOVA with Dunnett’s T3 multiple comparisons test.

## DISCUSSION

There remain many gaps in our knowledge of the diverse ways that viruses interact with and modulate host cells, which can include direct interactions with cellular miRNAs. Here we adapted CLEAR-CLIP as a powerful method to investigate genome-wide analysis of miRNA-target interactions in Mock– and RSV-infected A549 lung epithelial cells. After initial optimisation, we were able to recover a high number of miRNA-target chimeras in both conditions which allowed us to identify viral as well as host gene targets in this lung cell line. We show for the first time that RSV interacts directly with miR-26 and miR-27 miRNA families. We do not rule out that other miRNAs could also interact with RSV to a degree and note multiple binding sites in the SH and G gene for let-7, which is a highly abundant miRNA family that has also been found to bind to *Pestiviruses* (4). However, our data highlight specificity in the binding of miR-26 and miR-27 where the chimeras are concentrated at individual seed-based sites in the genome. Further work is required to understand whether the interactions with the miRNAs directly influence the RSV life cycle. Here we have focused on defining the host target networks of these two miRNAs and examining the scope for de-repression of their targets upon infection.

Our analysis of HC target chimeras with miR-27 suggests an overall decrease in host target interactions upon infection with correlated up-regulation of specific targets, including CCNG1 and PHLPP2 that have been associated with inhibition of the cell cycle (57–61). The role of miR-27 in regulating (enhancing) cell cycle is well documented (79–83), as is the importance of cell cycle arrest for RSV infection, which has been shown in both A549 cells and primary human bronchial epithelial cells (84,85). We show that targets associated with cell cycle are the most sensitive to miR-27 inhibition in A549 cells, based on a ranked list of expression changes of targets in cells treated with a miR-27 inhibitor. Our working model from the chimeric and gene expression data is that RSV interaction with miR-27 causes a de-repression of certain targets in A549s and in particular those involved in cell cycle regulation. However, it is likely that miR-27 is not the sole regulator of the cell cycle in infection but rather acts in coordination with other factors. In addition, we note that additional genes from the list of 395 HC miR-27 targets that we identify here are likely consequential to RSV infection including those associated with metabolism (GOLM1 and LIMK1) and immune signalling (CSF1) which require further investigation *in vivo*. A previous study in MCMV found that viral mutants unable to degrade miR-27 showed a significant reduction in viral titre in the acute infection in the lung (8). We previously showed that miR-27 has antiviral properties in different respiratory viruses *in vitro*, including multiple influenza strains, RSV A2 and a more pronounced effect against RSV Bt2a, a recent clinical isolate of RSV (86).

Our study also identifies miR-26 as a miRNA that directly interacts with RSV. This miRNA has not been reported yet to interact with other viruses but our pathway analysis of the 165 HC target sites demonstrates integral link with the host response to the virus and antiviral immunity. In contrast to miR-27, globally we do not see a significant decrease in miR-26 HC chimeric reads upon infection and we describe an association of miR-26 with antiviral immunity through targeting a network of genes induced upon infection, for example OAS2 as shown in this study. One miR-26 target that does appear to be de-repressed upon infection is the DEAD-box helicase 3 X-linked, DDX3X, which has been shown to be required for virulence of 18 different viruses (56), including RSV (87) and is also important in the antiviral immune response. miR-26 has been reported to regulate and boost antiviral immunity in several different viral infections and has been proposed as a broad-spectrum antiviral (88–91). Further work is required to understand the precise role of the miR-26-RSV interaction *in vivo*.

As exemplified in our study, a challenge in validating the mechanism of action of a miRNA is the sheer number of targets that (collectively) may underpin its function. However, another under-appreciated challenge in studying miRNA functions is the gap in knowledge on the non-3’UTR targets they may also regulate. The most up-to-date prediction programs (targetscan.org) and validated target databases (miRTarBase) are focused on 3’UTR or CDS targets. Yet a burgeoning body of data suggest miRNAs can also have nuclear functions. Using our method to select for high confidence target sites, we examined whether there is support for other target classes in our dataset. Beyond 3’UTRs and CDS we show that pseudogenes (PSGs) and lncRNAs overlapping regulatory regions are strong candidates for functional miRNA interactions. We identify a lncRNA with a highly conserved miR-27 site which is derived from a pseudogene recently described to be expressed downstream of CEACAM1. We validate nuclear expression of miR-27 as well as the pseudogene and show that knockdown of the pseudogene influences CEACAM1 expression. Further work is required to understand whether and how the interaction of miR-27 and/or let-7 (which also binds to this lncRNA) regulates transcription. Our work brings biochemical evidence for targets of miR-26 and miR-27 that are directly implicated in RSV infection, including the virus itself and also interfaces with a growing body of literature demonstrating that targets beyond those mapping to the 3’UTR should be considered when trying to understand miRNA and AGO2 functions in infection and disease (13,92).

## Data Availability

CLEAR-CLIP raw files as well as a processed data table containing all human miRNA-host target interactions have been deposited in the Gene Expression Omnibus (GEO) under accession number GSE232686 (https://www.ncbi.nlm.nih.gov/geo/query/acc.cgi?acc=GSE232686). Total RNA-seq of Mock and RSV-infected A549 as well as A549 WT, A549 miR-27a and miR-27 mimics/inhibitors have been deposited in GEO under accession numbers GSE231788 (https://www.ncbi.nlm.nih.gov/geo/query/acc.cgi?acc=GSE231788) and GSE231787 (https://www.ncbi.nlm.nih.gov/geo/query/acc.cgi?acc=GSE231787), respectively. The small RNA-seq data of Mock and RSV-infected A549 has been deposited in GEO under accession number GSE231784 (https://www.ncbi.nlm.nih.gov/geo/query/acc.cgi?acc=GSE231784). The original code used for the CLEAR-CLIP analysis is available at https://doi.org/10.5281/zenodo.7924850

## Acknowledgements

The authors thank Dr Alfonso Garrido-Lecca from the Yi group at the University of Boulder for sharing advice on modifications to the CLEAR-CLIP protocol. The authors thank the Macias group from the University of Edinburgh for sharing RT-qPCR primers, as well as the IgG control antibody. The authors also thank Dr Pablo Tristán-Ramos from the Heras group at the University of Granada for discussions around crosslinking conditions during the CLEAR-CLIP protocol optimisation and Dr Anton McCaffrey and Sabrina Shore from Trilink for discussions around the modified 3’ adapter. Total RNA-seq library preparation and sequencing was carried out by the Genetics core at the Edinburgh Clinical Research Facility. Small RNA sequencing was carried out by Edinburgh Genomics at the University of Edinburgh.

## Author Contributions

S.R. and A.H.B designed the study. All authors participated in the discussion and interpretation of the results. S.R. and J.L. performed the experiments. K.G. optimised the CLEAR-CLIP protocol for the fluorescent 3’ adapter. J.W. provided the virus and reagents. S.K. developed the CLEAR-CLIP bioinformatic pipeline in communication with S.R. and C.A.G. J.R.B.B. modified the CLEAR-CLIP pipeline. S.K., J.R.B.B., SR performed the bioinformatic analysis of the (small) RNA-seq data. J.R.B.B and S.R. analysed the data with input from A.H.B and C.A.G. S.R. and A.H.B. wrote the manuscript. All authors revised the manuscript.

---

**Figure S1.**
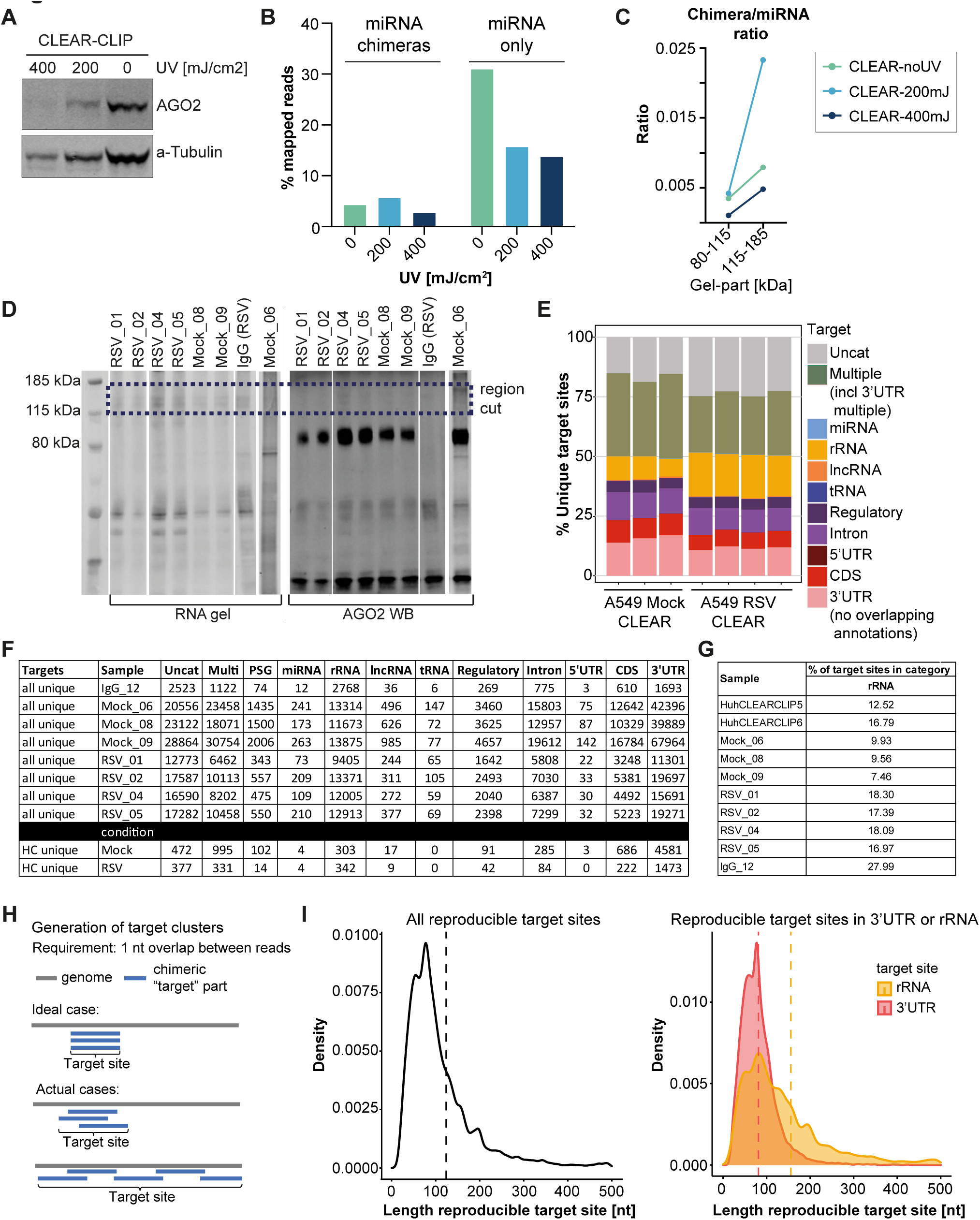
CLEAR-CLIP protocol optimisations and target site annotations and filtering. (**A**) Western blot analysis of protein lysates after crosslinking A549 cells with different doses of UV, probed for AGO2 and α–Tubulin (loading control). Equivalent of 5 x 10^5^ cells loaded per lane. (**B**) Percentage of chimeric reads and miRNA only reads mapped to the human genome for different UV crosslinking dosages. (**C**) Ratio of chimeric reads compared to miRNA only reads of two different molecular weight ranges purified from SDS-PAGE (80-115 kDa vs 115-185 kDa), separated by different crosslinking conditions. (**D**) Visualisation of the infrared 3’ adapter ligated RNA (left) and AGO2 protein (right) on the SDS-PAGE before purifying complexes from 115-185 kDa. Western blot (WB) for AGO2 was done separately to the RNA gel with 8 µl of the total eluate. (**E**) Percentage of unique target sites across different RNA categories, similar to Figure 1E. For this graph 3’UTRs that were overlapping other annotations were classified as Multiple and the 3’UTR category only contains target sites that exclusively map to 3’UTRs. (**F**) Table showing the number of unique target sites across different RNA categorises that were used to calculate the percentages for the Figures 1E & 1F. (**G**) Table showing the percentage of target sites mapped to rRNA across different samples. HuhCLEARCLIP5/6 are from Moore et al. data (16). (**H**) Schematic of the generation of target site clusters. Visualising that the required minimum overlap of 1 nt can result in extensive target sites with only little overlap between reads. (**I**) Density plot showing the length distribution of target sites in nt. Dashed line indicates mean target site length. Left panel: all reproducible target sites; right panel: comparing target sites from 3’UTRs to rRNA target sites.

**Figure S2.**
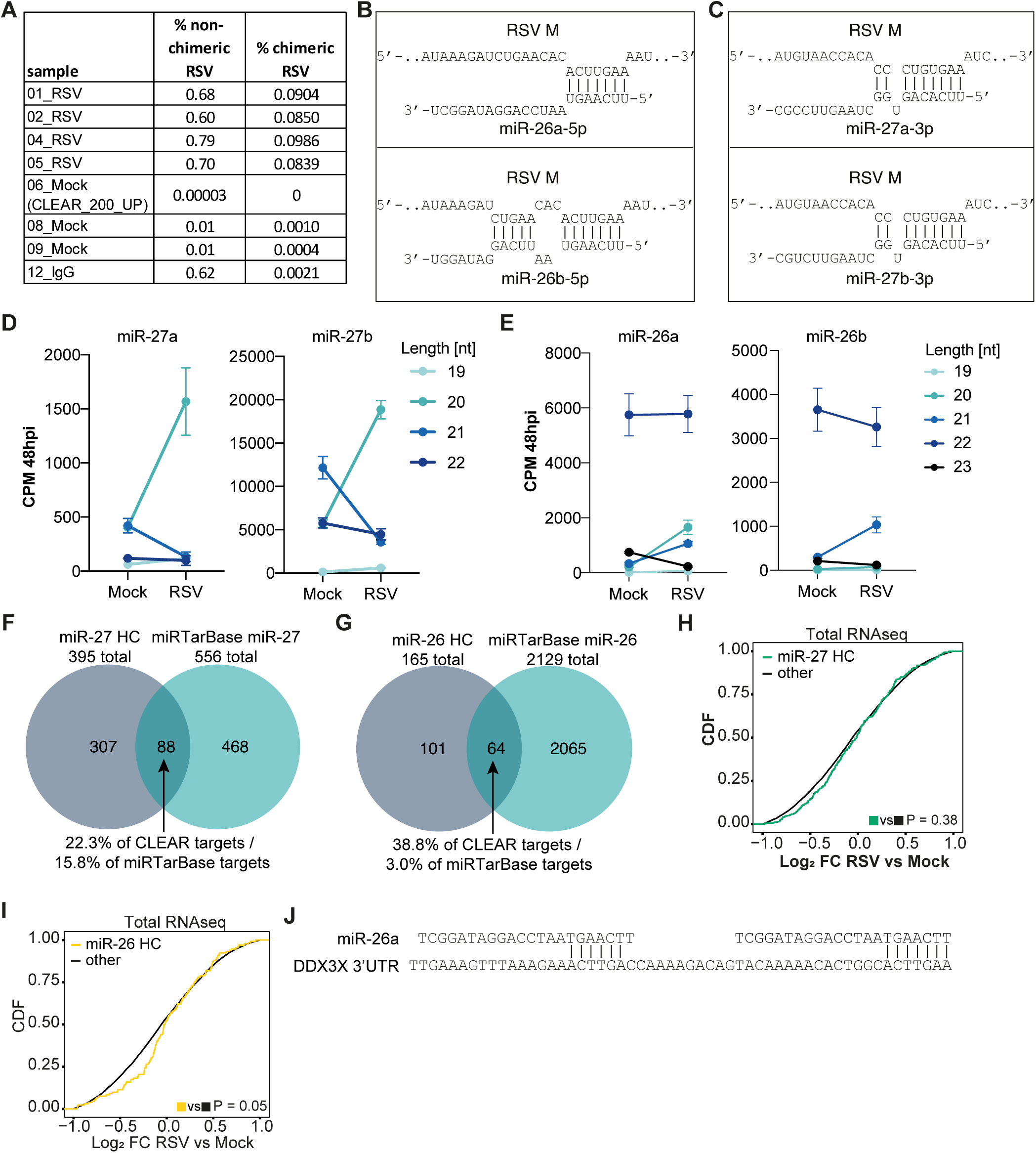
miRNA-RSV interactions, miR-26 and miR-27 miRNA length and HC targets in RSV-infection. (**A**) Table showing percentage of non-chimeric and chimeric reads mapping to RSV genome across the different samples. (**B-C**) Predicted binding pattern of miR-26 (**B**) and miR-27 (**C**) family members for the most dominant target site in the RSV genome, determined with IntaRNA (35). (**D-E**) miRNA length in nt in Mock and RSV-infected samples at 48 hpi, determined from smallRNA-seq data for miR-27 (**D**) and miR-26 (**E**). Data is shown as mean ± SD of three biological replicates per condition. (**F-G**) Venn diagram comparing the overlap of HC target interactions found in our CLEAR-CLIP data compared to validated targets in miRTarBase (49) for miR-27 (**F**) and miR-26 (**G**). (**H, I**) CDF plot of total RNA-seq data showing log_2_ fold change in expression of miRNA HC 3’UTR target genes vs non-target genes in RSV vs Mock-infected cells for miR-27 (**H**) and miR-26 (**I**). Two-tailed Wilcoxon p-values are shown for the comparisons. (**J**) Schematic of miR-26 seed sites within the target site in DDX3X 3’UTR.

**Figure S3.**
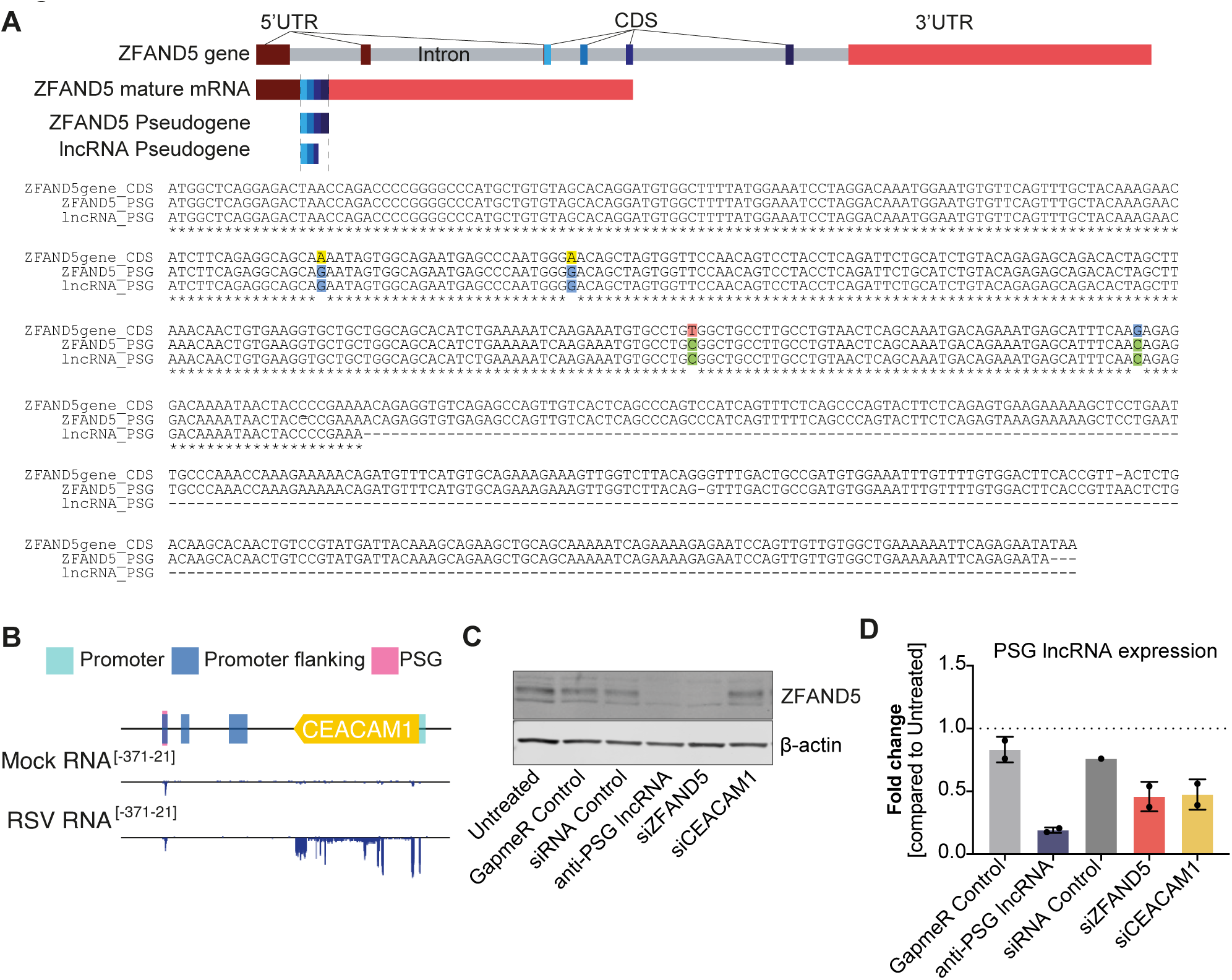
ZFAND5 pseudogene lncRNA and ZFAND5 parental gene characterisation and expression upon knockdown. (**A**)Schematic of the parental ZFAND5 gene locus, as well as mature ZFAND5 mRNA, ZFAND5 PSG, and PSG lncRNA (top). Clustal Omega multiple sequence alignment if ZFAND5 CDS, ZFAND5 PSG, and PSG lncRNA (bottom). Nucleotide differences are highlighted. (**B**) Schematic of CEACAM1 gene locus in comparison to PSG, showing below expression of CEACAM1 and PSG in Mock and RSV determined from total RNA-seq. (**C**) Western blot analysis of ZFAND5 protein expression in A549 cells after transfection with 5 nM GapmeR (anti-PSG lncRNA, GapmeR Control) or 10 nM of siRNAs and siRNA control compared to β-Actin loading control. (**D**) RT-qPCR results of PSG lncRNA expression after transfection with 5 nM GapmeR or 10 nM siRNAs. Data is shown as mean ± SD for two biological replicates.

## Supplemental Tables

**Table S1**: Summary of CLEAR-CLIP data sets generated in this study, as well as CLEARCLIP samples generated by Moore et al. Information for each sample includes sample name, UV intensity used for crosslinking (UV), molecular weight of protein-RNA complexes purified (MW region), information about sequencing run, as well as information about non-chimeric and chimeric reads with both the human and RSV genome. ND = not determined.

**Table S2**: miRNA chimeric reads with RSV genome (mirfirst = interacting miRNA; mirfirstclustercoord = miRNA name together with RSV genome coordinates of the target site; RSV_01 – IgG_12 = read counts of target site in the different samples; HC = reproducible, seed-based target sites, 1 = HC site, 0 = no HC site; sum reads = sum of reads in RSV_01, RSV_02, RSV_04, RSV_05).

**Table S3**: HC target sites for miR-26 and miR-27 with the human genome (Chr/start/end = genome location of target site; mirfirst = interacting miRNA; RSV_01 – IgG_12 = read counts of target site in the different samples; strand = indicates positive (+) or negative strand (-); UTR3 – unprocessed_pseudogene = gene symbol/Ensembl ID of genome annotations in the target site location; targetseq = sequence of target site).

**Table S4**: Differentially expressed genes in RSV vs Mock-infected cells at 48 hpi, MOI of 0.1 (log2FC = log_2_ fold change, FDR = False discovery rate, miR-27ab_HC = high confidence target of miR-27a/b with 1 = miR-27 HC target & 0 = non-miR-27 HC target).

**Table S5**: ShinyGO Biological process pathway analysis of miR-27 HC 3’UTR targets (nGenes = number of genes).

**Table S6**: ShinyGO Biological process pathway analysis of miR-26 HC 3’UTR targets (nGenes = number of genes).

**Table S7**: Differential gene expression in response to miR-27 mimics/inhibitors transfection compared to controls (logFC = log_2_ fold change, FDR = False discovery rate, miR-27HC = high confidence target of miR-27a/b with 1 = miR-27 HC target & NA = non-miR-27 HC target).

**Table S8**: HC target sites in regulatory regions overlapping either lncRNAs or pseudogenes (Chr/start/end = genome location of target site; mirfirst = interacting miRNA; RSV_01 – IgG_12 = read counts of target site in the different samples; strand = indicates positive (+) or negative strand (-); lncRNA – unprocessed_pseudogene = gene symbol/Ensembl ID of genome annotations in target site location; targetseq = sequence of target site).

**Table S9**: Oligonucleotides, miRNA mimics & inhibitors, siRNAs and GapmeRs used in this study.

## REFERENCES

1. Jopling, C.L., Yi, M., Lancaster, A.M., Lemon, S.M. and Sarnow, P. (2005) Modulation of Hepatitis C Virus RNA Abundance by a Liver-Specific MicroRNA. Science, 309, 1577–1581.

2. Shimakami, T., Yamane, D., Jangra, R.K., Kempf, B.J., Spaniel, C., Barton, D.J. and Lemon, S.M. (2012) Stabilization of hepatitis C virus RNA by an Ago2-miR-122 complex. Proc Natl Acad Sci U S A, 109, 941–946.

3. Luna, J.M., Scheel, T.K., Danino, T., Shaw, K.S., Mele, A., Fak, J.J., Nishiuchi, E., Takacs, C.N., Catanese, M.T., de Jong, Y.P., et al. (2015) Hepatitis C virus RNA functionally sequesters miR-122. Cell, 160, 1099–1110.

4. Scheel, T.K., Luna, J.M., Liniger, M., Nishiuchi, E., Rozen-Gagnon, K., Shlomai, A., Auray, G., Gerber, M., Fak, J., Keller, I. et al. (2016) A Broad RNA Virus Survey Reveals Both miRNA Dependence and Functional Sequestration. Cell Host Microbe, 19, 409–423.

5. Cazalla, D., Yario, T. and Steitz, J.A. (2010) Down-regulation of a host microRNA by a Herpesvirus saimiri noncoding RNA. Science, 328, 1563–1566.

6. Buck, A.H., Perot, J., Chisholm, M.A., Kumar, D.S., Tuddenham, L., Cognat, V., Marcinowski, L., Dolken, L. and Pfeffer, S. (2010) Post-transcriptional regulation of miR-27 in murine cytomegalovirus infection. RNA, 16, 307–315.

7. Libri, V., Helwak, A., Miesen, P., Santhakumar, D., Borger, J.G., Kudla, G., Grey, F., Tollervey, D. and Buck, A.H. (2012) Murine cytomegalovirus encodes a miR-27 inhibitor disguised as a target. Proc Natl Acad Sci U S A, 109, 279–284.

8. Marcinowski, L., Tanguy, M., Krmpotic, A., Radle, B., Lisnic, V.J., Tuddenham, L., Chane-Woon-Ming, B., Ruzsics, Z., Erhard, F., Benkartek, C. et al. (2012) Degradation of cellular mir-27 by a novel, highly abundant viral transcript is important for efficient virus replication in vivo. PLoS Pathog, 8, e1002510.

9. Janssen, H.L.A., Reesink, H.W., Lawitz, E.J., Zeuzem, S., Rodriguez-Torres, M., Patel, K., van der Meer, A.J., Patick, A.K., Chen, A., Zhou, Y., et al. (2013) Treatment of HCV Infection by Targeting MicroRNA. New England Journal of Medicine, 368, 1685–1694.

10. van der Ree, M.H., de Vree, J.M., Stelma, F., Willemse, S., van der Valk, M., Rietdijk, S., Molenkamp, R., Schinkel, J., van Nuenen, A.C., Beuers, U., et al. (2017) Safety, tolerability, and antiviral effect of RG-101 in patients with chronic hepatitis C: a phase 1B, double-blind, randomised controlled trial. The Lancet, 389, 709–717.

11. Agarwal, V., Bell, G.W., Nam, J.W. and Bartel, D.P. (2015) Predicting effective microRNA target sites in mammalian mRNAs. Elife, 4.

12. Chu, Y., Yokota, S., Liu, J., Kilikevicius, A., Johnson, K.C. and Corey, D.R. (2021) Argonaute binding within human nuclear RNA and its impact on alternative splicing. Rna, 27, 991–1003.

13. Stavast, C.J. and Erkeland, S.J. (2019) The Non-Canonical Aspects of MicroRNAs: Many Roads to Gene Regulation. Cells, 8.

14. Helwak, A., Kudla, G., Dudnakova, T. and Tollervey, D. (2013) Mapping the human miRNA interactome by CLASH reveals frequent noncanonical binding. Cell, 153, 654–665.

15. Grosswendt, S., Filipchyk, A., Manzano, M., Klironomos, F., Schilling, M., Herzog, M., Gottwein, E. and Rajewsky, N. (2014) Unambiguous identification of miRNA:target site interactions by different types of ligation reactions. Mol Cell, 54, 1042–1054.

16. Moore, M.J., Scheel, T.K., Luna, J.M., Park, C.Y., Fak, J.J., Nishiuchi, E., Rice, C.M. and Darnell, R.B. (2015) miRNA-target chimeras reveal miRNA 3’-end pairing as a major determinant of Argonaute target specificity. Nat Commun, 6, 8864.

17. Mammas, I.N., Drysdale, S.B., Rath, B., Theodoridou, M., Papaioannou, G., Papatheodoropoulou, A., Koutsounaki, E., Koutsaftiki, C., Kozanidou, E., Achtsidis, V. et al. (2020) Update on current views and advances on RSV infection (Review). Int J Mol Med, 46, 509–520.

18. Hallak, L.K., Spillmann, D., Collins, P.L. and Peeples, M.E. (2000) Glycosaminoglycan Sulfation Requirements for Respiratory Syncytial Virus Infection. Journal of Virology, 74, 10508.

19. Guerrero-Plata, A., Casola, A., Suarez, G., Yu, X., Spetch, L., Peeples, M.E. and Garofalo, R.P. (2006) Differential Response of Dendritic Cells to Human Metapneumovirus and Respiratory Syncytial Virus. American Journal of Respiratory Cell and Molecular Biology, 34, 320–329.

20. Bjerke, G.A. and Yi, R. (2020) Integrated analysis of directly captured microRNA targets reveals the impact of microRNAs on mammalian transcriptome. Rna, 26, 306–323.

21. Kwok, S., Kellogg, D.E., McKinney, N., Spasic, D., Goda, L., Levenson, C. and Sninsky, J.J. (1990) Effects of primer-template mismatches on the polymerase chain reaction: Human immunodeficiency virus type 1 model studies. Nucleic Acids Research, 18, 999–1005.

22. Bru, D., Martin-Laurent, F. and Philippot, L. (2008) Quantification of the Detrimental Effect of a Single Primer-Template Mismatch by Real-Time PCR Using the 16S rRNA Gene as an Example. Applied and Environmental Microbiology, 74, 1660–1663.

23. Stadhouders, R., Pas, S.D., Anber, J., Voermans, J., Mes, T.H.M. and Schutten, M. (2010) The Effect of Primer-Template Mismatches on the Detection and Quantification of Nucleic Acids Using the 5′ Nuclease Assay. The Journal of Molecular Diagnostics, 12, 109–117.

24. Livak, K.J. and Schmittgen, T.D. (2001) Analysis of relative gene expression data using real-time quantitative PCR and the 2(-Delta Delta C(T)) Method. Methods, 25, 402–408.

25. Burke, J.M. and Sullivan, C.S. (2017) DUSP11 – An RNA phosphatase that regulates host and viral non-coding RNAs in mammalian cells. RNA Biology, 14, 1457–1465.

26. Gagnon, Keith T., Li, L., Chu, Y., Janowski, Bethany A. and Corey, David R. (2014) RNAi Factors Are Present and Active in Human Cell Nuclei. Cell Reports, 6, 211–221.

27. Schindelin, J., Arganda-Carreras, I., Frise, E., Kaynig, V., Longair, M., Pietzsch, T., Preibisch, S., Rueden, C., Saalfeld, S., Schmid, B. et al. (2012) Fiji: an open-source platform for biological-image analysis. Nature Methods, 9, 676–682.

28. Stael, S., Miller, L.P., Fernández-Fernández, Á.D. and Van Breusegem, F. (2022) In Klemenčič, M., Stael, S. and Huesgen, P. F. (eds.), Plant Proteases and Plant Cell Death: Methods and Protocols. Springer US, New York, NY, pp. 127–137.

29. Pall, G.S. and Hamilton, A.J. (2008) Improved northern blot method for enhanced detection of small RNA. Nat Protoc, 3, 1077–1084.

30. Martin, M. (2011) Cutadapt removes adapter sequences from high-throughput sequencing reads. 2011, 17, 3.

31. Zhao, S., Gordon, W., Du, S., Zhang, C., He, W., Xi, L., Mathur, S., Agostino, M., Paradis, T., von Schack, D., et al. (2017) QuickMIRSeq: a pipeline for quick and accurate quantification of both known miRNAs and isomiRs by jointly processing multiple samples from microRNA sequencing. BMC Bioinformatics, 18, 180.

32. Langmead, B. and Salzberg, S.L. (2012) Fast gapped-read alignment with Bowtie 2. Nature Methods, 9, 357–359.

33. Axtell, M.J. (2013) ShortStack: comprehensive annotation and quantification of small RNA genes. RNA, 19, 740–751.

34. Quinlan, A.R. and Hall, I.M. (2010) BEDTools: a flexible suite of utilities for comparing genomic features. Bioinformatics, 26, 841–842.

35. Mann, M., Wright, P.R. and Backofen, R. (2017) IntaRNA 2.0: enhanced and customizable prediction of RNA–RNA interactions. Nucleic Acids Research, 45, W435–W439.

36. Lawrence, M., Huber, W., Pagès, H., Aboyoun, P., Carlson, M., Gentleman, R., Morgan, M.T. and Carey, V.J. (2013) Software for Computing and Annotating Genomic Ranges. PLOS Computational Biology, 9, e1003118.

37. Lawrence, M., Gentleman, R. and Carey, V. (2009) rtracklayer: an R package for interfacing with genome browsers. Bioinformatics, 25, 1841–1842.

38. Robinson, J.T., Thorvaldsdottir, H., Winckler, W., Guttman, M., Lander, E.S., Getz, G. and Mesirov, J.P. (2011) Integrative genomics viewer. Nat Biotechnol, 29, 24–26.

39. Bolger, A.M., Lohse, M. and Usadel, B. (2014) Trimmomatic: a flexible trimmer for Illumina sequence data. Bioinformatics, 30, 2114–2120.

40. Kim, D., Paggi, J.M., Park, C., Bennett, C. and Salzberg, S.L. (2019) Graph-based genome alignment and genotyping with HISAT2 and HISAT-genotype. Nature Biotechnology, 37, 907–915.

41. Liao, Y., Smyth, G.K. and Shi, W. (2019) The R package Rsubread is easier, faster, cheaper and better for alignment and quantification of RNA sequencing reads. Nucleic Acids Research, 47, e47–e47.

42. Fu, X., Liu, P., Dimopoulos, G. and Zhu, J. (2020) Dynamic miRNA-mRNA interactions coordinate gene expression in adult Anopheles gambiae. PLoS Genet, 16, e1008765.

43. Stebel, S., Breuer, J. and Rossbach, O. (2022) Studying miRNA-mRNA Interactions: An Optimized CLIP-Protocol for Endogenous Ago2-Protein. Methods and Protocols, 10.3390/mps5060096.

44. Chi, S.W., Zang, J.B., Mele, A. and Darnell, R.B. (2009) Argonaute HITS-CLIP decodes microRNA-mRNA interaction maps. Nature, 460, 479–486.

45. Chandradoss, Stanley D., Schirle, Nicole T., Szczepaniak, M., MacRae, Ian J. and Joo, C. (2015) A Dynamic Search Process Underlies MicroRNA Targeting. Cell, 162, 96–107.

46. Bartel, D.P. (2018) Metazoan MicroRNAs. Cell, 173, 20–51.

47. McCaskill, J., Praihirunkit, P., Sharp, P.M. and Buck, A.H. (2015) RNA-mediated degradation of microRNAs: A widespread viral strategy? RNA Biol, 12, 579–585.

48. Haas, G., Cetin, S., Messmer, M., Chane-Woon-Ming, B., Terenzi, O., Chicher, J., Kuhn, L., Hammann, P. and Pfeffer, S. (2016) Identification of factors involved in target RNA-directed microRNA degradation. Nucleic Acids Research, 44, 2873–2887.

49. Tomasello, L., Distefano, R., Nigita, G. and Croce, C.M. (2021) The MicroRNA Family Gets Wider: The IsomiRs Classification and Role. Frontiers in Cell and Developmental Biology, 9.

50. Huang, H.-Y., Lin, Y.-C.-D., Cui, S., Huang, Y., Tang, Y., Xu, J., Bao, J., Li, Y., Wen, J., Zuo, H. et al. (2022) miRTarBase update 2022: an informative resource for experimentally validated miRNA–target interactions. Nucleic Acids Research, 50, D222–D230.

51. Ge, S.X., Jung, D. and Yao, R. (2019) ShinyGO: a graphical gene-set enrichment tool for animals and plants. Bioinformatics, 36, 2628–2629.

52. Altan-Bonnet, G. and Mukherjee, R. (2019) Cytokine-mediated communication: a quantitative appraisal of immune complexity. Nature Reviews Immunology, 19, 205–217.

53. Choi, U.Y., Kang, J.-S., Hwang, Y.S. and Kim, Y.-J. (2015) Oligoadenylate synthase-like (OASL) proteins: dual functions and associations with diseases. Experimental & Molecular Medicine, 47, e144–e144.

54. Forte, E. and Luftig, M.A. (2009) MDM2-dependent inhibition of P53 is required for Epstein-Barr virus B cell growth transformation and infected cell survival. Infectious Agents and Cancer, 4, P26.

55. Muñoz-Fontela, C., Mandinova, A., Aaronson, S.A. and Lee, S.W. (2016) Emerging roles of p53 and other tumour-suppressor genes in immune regulation. Nature Reviews Immunology, 16, 741–750.

56. Winnard, P.T., Vesuna, F. and Raman, V. (2021) Targeting host DEAD-box RNA helicase DDX3X for treating viral infections. Antiviral Research, 185, 104994.

57. Kimura, S.H., Ikawa, M., Ito, A., Okabe, M. and Nojima, H. (2001) Cyclin G1 is involved in G2/M arrest in response to DNA damage and in growth control after damage recovery. Oncogene, 20, 3290–3300.

58. Taguchi, K., Elias, B.C., Sugahara, S., Sant, S., Freedman, B.S., Waikar, S.S., Pozzi, A., Zent, R., Harris, R.C., Parikh, S.M. et al. (2022) Cyclin G1 induces maladaptive proximal tubule cell dedifferentiation and renal fibrosis through CDK5 activation. J Clin Invest, 132.

59. Lin, T., Ma, Q., Zhang, Y., Zhang, H., Yan, J. and Gao, C. (2018) MicroRNA-27a functions as an oncogene in human osteosarcoma by targeting CCNG1. Oncol Lett, 15, 1067–1071.

60. Brognard, J., Sierecki, E., Gao, T. and Newton, A.C. (2007) PHLPP and a Second Isoform, PHLPP2, Differentially Attenuate the Amplitude of Akt Signaling by Regulating Distinct Akt Isoforms. Molecular Cell, 25, 917–931.

61. Xian, Q., Zhao, R. and Fu, J. (2020) MicroRNA-527 Induces Proliferation and Cell Cycle in Esophageal Squamous Cell Carcinoma Cells by Repressing PH Domain Leucine-Rich-Repeats Protein Phosphatase 2. Dose-Response, 18, 1559325820928687.

62. Bongolo, C.C., Thokerunga, E., Yan, Q., Yacouba, M.B.M. and Wang, C. (2022) Exosomes Derived from microRNA-27a-3p Overexpressing Mesenchymal Stem Cells Inhibit the Progression of Liver Cancer through Suppression of Golgi Membrane Protein 1. Stem Cells International, 2022, 9748714.

63. Frans, M.T., Kuipers, E.M., Bianchi, F. and van den Bogaart, G. (2023) Unveiling the impact of GOLM1/GP73 on cytokine production in cancer and infectious disease. Immunology & Cell Biology, 101, 727–734.

64. Nagaraj, M., Höring, M., Ahonen, M.A., Nguyen, V.D., Zhou, Y., Vihinen, H., Jokitalo, E., Liebisch, G., Nidhina Haridas, P.A. and Olkkonen, V.M. (2022) GOLM1 depletion modifies cellular sphingolipid metabolism and adversely affects cell growth. Journal of Lipid Research, 63.

65. Avota, E., Bodem, J., Chithelen, J., Mandasari, P., Beyersdorf, N. and Schneider-Schaulies, J. (2021) The Manifold Roles of Sphingolipids in Viral Infections. Front Physiol, 12, 715527.

66. Lu, Y., Xu, S., Sun, H., Shan, J., Shen, C., Ji, J., Lin, L., Xu, J., Peng, L., Dai, C. et al. (2023) Analysis of temporal metabolic rewiring for human respiratory syncytial virus infection by integrating metabolomics and proteomics. Metabolomics, 19, 30.

67. Wen, X., Ding, L., Wang, J.-J., Qi, M., Hammonds, J., Chu, H., Chen, X., Hunter, E. and Spearman, P. (2014) ROCK1 and LIM Kinase Modulate Retrovirus Particle Release and Cell-Cell Transmission Events. Journal of Virology, 88, 6906–6921.

68. Vorster, P.J., Guo, J., Yoder, A., Wang, W., Zheng, Y., Xu, X., Yu, D., Spear, M. and Wu, Y. (2011) LIM Kinase 1 Modulates Cortical Actin and CXCR4 Cycling and Is Activated by HIV-1 to Initiate Viral Infection *. Journal of Biological Chemistry, 286, 12554–12564.

69. Xiang, Y., Zheng, K., Zhong, M., Chen, J., Wang, X., Wang, Q., Wang, S., Ren, Z., Fan, J. and Wang, Y. (2014) Ubiquitin-proteasome-dependent slingshot 1 downregulation in neuronal cells inactivates cofilin to facilitate HSV-1 replication. Virology, 449, 88–95.

70. König, R., Stertz, S., Zhou, Y., Inoue, A., Hoffmann, H.H., Bhattacharyya, S., Alamares, J.G., Tscherne, D.M., Ortigoza, M.B., Liang, Y. et al. (2010) Human host factors required for influenza virus replication. Nature, 463, 813–817.

71. Villalonga, E., Mosrin, C., Normand, T., Girardin, C., Serrano, A., Žunar, B., Doudeau, M., Godin, F., Bénédetti, H. and Vallée, B. (2023) LIM Kinases, LIMK1 and LIMK2, Are Crucial Node Actors of the Cell Fate: Molecular to Pathological Features. Cells, 12, 805.

72. Yi, F., Guo, J., Dabbagh, D., Spear, M., He, S., Kehn-Hall, K., Fontenot, J., Yin, Y., Bibian, M., Park Chul, M. et al. (2017) Discovery of Novel Small-Molecule Inhibitors of LIM Domain Kinase for Inhibiting HIV-1. Journal of Virology, 91, 10.1128/jvi.02418-02416.

73. Benhamed, M., Herbig, U., Ye, T., Dejean, A. and Bischof, O. (2012) Senescence is an endogenous trigger for microRNA-directed transcriptional gene silencing in human cells. Nat Cell Biol, 14, 266–275.

74. Matsui, M., Chu, Y., Zhang, H., Gagnon, K.T., Shaikh, S., Kuchimanchi, S., Manoharan, M., Corey, D.R. and Janowski, B.A. (2013) Promoter RNA links transcriptional regulation of inflammatory pathway genes. Nucleic Acids Res, 41, 10086–10109.

75. Di Mauro, V., Crasto, S., Colombo, F.S., Di Pasquale, E. and Catalucci, D. (2019) Wnt signalling mediates miR-133a nuclear re-localization for the transcriptional control of Dnmt3b in cardiac cells. Sci Rep, 9, 9320.

76. Turunen, T.A., Roberts, T.C., Laitinen, P., Vaananen, M.A., Korhonen, P., Malm, T., Yla-Herttuala, S. and Turunen, M.P. (2019) Changes in nuclear and cytoplasmic microRNA distribution in response to hypoxic stress. Sci Rep, 9, 10332.

77. Laitinen, P., Väänänen, M.-A., Kolari, I.-L., Mäkinen, P.I., Kaikkonen, M.U., Weinberg, M.S., Morris, K.V., Korhonen, P., Malm, T., Ylä-Herttuala, S. et al. (2022) Nuclear microRNA-466c regulates Vegfa expression in response to hypoxia. PLOS ONE, 17, e0265948.

78. Staszak, K. and Makałowska, I. (2021) Cancer, Retrogenes, and Evolution. Life, 11, 72.

79. Tian, Y., Fu, S., Qiu, G.-B., Xu, Z.-M., Liu, N., Zhang, X.-W., Chen, S., Wang, Y., Sun, K.-L. and Fu, W.-N. (2014) MicroRNA-27a promotes proliferation and suppresses apoptosis by targeting PLK2in laryngeal carcinoma. BMC Cancer, 14, 678.

80. Mertens-Talcott, S.U., Chintharlapalli, S., Li, X. and Safe, S. (2007) The Oncogenic microRNA-27a Targets Genes That Regulate Specificity Protein Transcription Factors and the G2-M Checkpoint in MDA-MB-231 Breast Cancer Cells. Cancer Research, 67, 11001–11011.

81. Lerner, M., Lundgren, J., Akhoondi, S., Jahn, A., Ng, H.-F., Moqadam, F.A., Oude Vrielink, J.A.F., Agami, R., Den Boer, M.L., Grandér, D., et al. (2011) MiRNA-27a controls FBW7/hCDC4-dependent cyclin E degradation and cell cycle progression. Cell Cycle, 10, 2172–2183.

82. Ye, P., Ke, X., Zang, X., Sun, H., Dong, Z., Lin, J., Wang, L., Liu, W., Miao, G., Tan, Y. et al. (2018) Up-regulated MiR-27-3p promotes the G1-S phase transition by targeting inhibitor of growth family member 5 in osteosarcoma. Biomedicine & Pharmacotherapy, 101, 219–227.

83. Su, C., Huang, D.P., Liu, J.W., Liu, W.Y. and Cao, Y.O. (2019) miR–27a–3p regulates proliferation and apoptosis of colon cancer cells by potentially targeting BTG1. Oncol Lett, 18, 2825–2834.

84. Gibbs, J.D., Ornoff, D.M., Igo, H.A., Zeng, J.Y. and Imani, F. (2009) Cell Cycle Arrest by Transforming Growth Factor β1 Enhances Replication of Respiratory Syncytial Virus in Lung Epithelial Cells. Journal of Virology, 83, 12424–12431.

85. Bian, T., Gibbs, J.D., Orvell, C. and Imani, F. (2012) Respiratory syncytial virus matrix protein induces lung epithelial cell cycle arrest through a p53 dependent pathway. PLoS One, 7, e38052.

86. McCaskill, J.L., Ressel, S., Alber, A., Redford, J., Power, U.F., Schwarze, J., Dutia, B.M. and Buck, A.H. (2017) Broad-Spectrum Inhibition of Respiratory Virus Infection by MicroRNA Mimics Targeting p38 MAPK Signaling. Mol Ther Nucleic Acids, 7, 256–266.

87. Yang, S.N.Y., Atkinson, S.C., Audsley, M.D., Heaton, S.M., Jans, D.A. and Borg, N.A. (2020) RK-33 Is a Broad-Spectrum Antiviral Agent That Targets DEAD-Box RNA Helicase DDX3X. Cells, 9, 170.

88. Zhao, F., Xu, G., Zhou, Y., Wang, L., Xie, J., Ren, S., Liu, S. and Zhu, Y. (2014) MicroRNA-26b Inhibits Hepatitis B Virus Transcription and Replication by Targeting the Host Factor CHORDC1 Protein*. Journal of Biological Chemistry, 289, 35029–35041.

89. Jia, X., Bi, Y., Li, J., Xie, Q., Yang, H. and Liu, W. (2015) Cellular microRNA miR-26a suppresses replication of porcine reproductive and respiratory syndrome virus by activating innate antiviral immunity. Sci Rep, 5, 10651.

90. Gao, S., Li, J., Song, L., Wu, J. and Huang, W. (2017) Influenza A virus-induced downregulation of miR-26a contributes to reduced IFNalpha/beta production. Virol Sin, 32, 261–270.

91. Zhang, J., Li, Z., Huang, J., Yin, H., Tian, J. and Qu, L. (2020) miR-26a Inhibits Feline Herpesvirus 1 Replication by Targeting SOCS5 and Promoting Type I Interferon Signaling. Viruses, 12, 2.

92. Chu, Y., Kilikevicius, A., Liu, J., Johnson, K.C., Yokota, S. and Corey, D.R. (2020) Argonaute binding within 3’-untranslated regions poorly predicts gene repression. Nucleic Acids Res, 48, 7439–7453.

